# Diversity and evolution of pigment types and the phycobilisome rod gene region of marine *Synechococcus* cyanobacteria

**DOI:** 10.1101/2021.06.21.449213

**Authors:** Théophile Grébert, Laurence Garczarek, Vincent Daubin, Florian Humily, Dominique Marie, Morgane Ratin, Alban Devailly, Gregory K. Farrant, Isabelle Mary, Daniella Mella-Flores, Gwen Tanguy, Karine Labadie, Patrick Wincker, David M. Kehoe, Frédéric Partensky

## Abstract

*Synechococcus* picocyanobacteria are ubiquitous and abundant photosynthetic organisms in the marine environment and contribute for an estimated 16% of the ocean net primary productivity. Their light-harvesting complexes, called phycobilisomes (PBS), are composed of a conserved allophycocyanin core from which radiates six to eight rods with variable phycobiliprotein and chromophore content. This variability allows *Synechococcus* to optimally exploit the wide variety of spectral niches existing in marine ecosystems. Seven distinct pigment types or subtypes have been identified so far in this taxon, based on the phycobiliprotein composition and/or the proportion of the different chromophores in PBS rods. Most genes involved in their biosynthesis and regulation are located in a dedicated genomic region called the PBS rod region. Here, we examined the variability of gene sequences and organization of this genomic region in a large set of sequenced isolates and natural populations of *Synechococcus* representative of all known pigment types. All regions start with a tRNA-Phe_GAA_ and some possess mobile elements including tyrosine recombinases, suggesting that their genomic plasticity relies on a tycheposon-like mechanism. Comparison of the phylogenies obtained for PBS and core genes revealed that the evolutionary history of PBS rod genes differs from the rest of the genome and is characterized by the co-existence of different alleles and frequent allelic exchange. We propose a scenario for the evolution of the different pigment types and highlight the importance of population-scale mechanisms in maintaining a wide diversity of pigment types in different *Synechococcus* lineages despite multiple speciation events.

## Introduction

As the second most abundant phytoplanktonic organism of the ocean, the picocyanobacterium *Synechococcus* plays a crucial role in the carbon cycle, accounting for about 16% of the ocean net primary productivity (Flombaum et al. 2013; Guidi et al. 2016). Members of this group are found from the equator to subpolar waters and from particle-rich river mouths to optically clear open ocean waters, environments displaying a wide range of nutrient concentrations, temperatures, light regimes and spectral niches (Olson et al. 1990; Wood et al. 1998; Farrant et al. 2016; Paulsen et al. 2016; Sohm et al. 2016; Grébert et al. 2018; Holtrop et al. 2021). This ecological success is tied to the remarkably large genetic (Ahlgren and Rocap 2012; Huang et al. 2012; Mazard et al. 2012; Farrant et al. 2016) and pigment diversity exhibited by these cells (Six et al. 2007; Humily et al. 2013; Grébert et al. 2018; Xia et al. 2018). The pigment diversity arises from wide variations in the composition of their light-harvesting antennae, called phycobilisomes (PBS).

The building blocks of PBS are phycobiliproteins, which consist of two subunits (α and β). Phycobiliproteins are assembled into trimers of heterodimers [(αβ)_3_] and then hexamers [(αβ)_6_] that are stacked into rod-like structures with the help of linker proteins (Yu and Glazer 1982; Adir 2005). Before PBS can be assembled into their typical fan-like structure, the α and β subunits must be modified by lyases that covalently attach one, two or three chromophores, called phycobilins, at conserved cysteine positions (Scheer and Zhao 2008; Schluchter et al. 2010; Bretaudeau et al. 2013). The PBS core is always made of allophycocyanin (APC), from which radiate six to eight rods (Wilbanks and Glazer 1993a; Sidler 1994). Three major pigment types have been defined thus far based on the phycobiliprotein composition of PBS rods (Six et al. 2007; Humily et al. 2013). The simplest rods are found in pigment type 1 (PT 1) and contain only phycocyanin (PC), which binds the red-light absorbing phycocyanobilin (PCB, *A*_max_ = 620-650 nm; Six et al. 2007). The rods of pigment type 2 (PT 2) contain PC and phycoerythrin-I (PE-I), which binds the green-light (GL) absorbing phycoerythrobilin (PEB; *A*_max_ = 545-560 nm). For pigment type 3 (PT 3), the rods contain the three types of phycobiliproteins, PC, PE-I and phycoerythrin-II (PE-II) and bind PCB, PEB and the blue-light (BL) absorbing phycourobilin (PUB, *A*_max_ = 495 nm; Ong et al. 1984; Six et al. 2007). Although these three phycobilins are isomers, PCB and PEB are created via oxidation/reduction reactions while PUB is generated by the isomerization of PEB during its covalent binding to a phycobiliprotein. This process is performed by dual-function enzymes called phycobilin lyase-isomerases (Scheer and Zhao 2008; Blot et al. 2009; Schluchter et al. 2010; Shukla et al. 2012; Sanfilippo, Nguyen, et al. 2019).

Five pigment subtypes have been further defined within PT 3, depending on their PUB:PEB ratio. This ratio is often approximated for living cells by the ratio of the PUB and PEB fluorescence excitation peaks at 495 and 545 nm (Exc_495:545_), with the emission measured at 585 nm. Subtype 3a strains have a fixed low Exc_495:545_ ratio (< 0.6) and are often called ‘GL specialists’, 3b strains have a fixed intermediate ratio (0.6 ≤ Exc_495:545_ < 1.6), while 3c strains display a fixed high Exc_495:545_ ratio (≥ 1.6) and are often called ‘BL specialists’ (Six et al. 2007). Strains belonging to subtype 3d dynamically tune their PUB:PEB ratio to the ambient GL:BL ratio, a reversible physiological process known as type IV chromatic acclimation (CA4; Palenik 2001; Everroad et al. 2006; Shukla et al. 2012; Humily et al. 2013; Sanfilippo et al. 2016; Sanfilippo, Nguyen, et al. 2019; Sanfilippo, Garczarek, et al. 2019). As a result, the Exc_495:545_ of these strains varies from 0.6 in GL to 1.6 in BL. Finally, the rare subtype 3e shows only faint changes in Exc_495:545_ when shifted between GL and BL (Humily et al. 2013).

Comparative genomics analysis of the first 11 marine *Synechococcus* sequenced genomes (Six et al. 2007) revealed that most genes encoding proteins involved in the biosynthesis and regulation of PBS rods are grouped into a single genomic location called the ‘PBS rod region’. These authors suggested that the gene content and organization of this region was specific of the different pigment types or subtypes, but they were unable to examine the degree of genomic and genetic variability for strains within each pigment type. Further sequencing of additional PT 3d strain genomes revealed that CA4 capability is correlated with the presence of a small genomic island that exists in one of two configurations, CA4-A and CA4-B, defining the two pigment genotypes 3dA and 3dB (Humily et al. 2013). A novel organization of the PBS rod region was also discovered, first from metagenomes from the Baltic Sea (Larsson et al. 2014) and then from strains isolated from the Black Sea (Sánchez-Baracaldo et al. 2019). Gene content analysis identified these as a new PT 2 genotype named PT 2B, while the original PT 2 was renamed PT 2A. Finally, the genome sequencing of the high-PUB-containing strains KORDI-100 and CC9616 showed that although they display a high, PT 3c-like Exc_495:545_ ratio, the gene complement, order, and alleles of their PBS rod region differ from PT 3c, establishing an additional pigment subtype called PT 3f (Mahmoud et al. 2017; Grébert et al. 2018; Xia et al. 2018). An interesting feature of the genes within the PBS rod region is that their evolutionary history apparently differs from that of the core genome (Six et al. 2007; Everroad and Wood 2012; Humily et al. 2014; Grébert et al. 2018; Carrigee, Frick, et al. 2020), but the reason(s) for this autonomy remains unclear.

Here, we provide a more comprehensive understanding of the phylogenetic and genomic diversity of *Synechococcus* pigment types by leveraging the large number of recently available genomes of marine *Synechococcus* and *Cyanobium* isolates for further comparative genomic analysis. We also use a targeted metagenomics approach to directly retrieve and analyze PBS rod regions from natural populations as well as single-cell amplified genomes (SAGs). This broadened exploration leads us to propose hypotheses for the evolution of the PBS rod regions as well as for the maintenance of the wide pigment diversity found in most lineages, despite multiple speciation events. This study highlights the importance of population-scale mechanisms such as lateral transfers and incomplete lineage sorting in shaping the distribution of pigment types among *Synechococcus* lineages.

## Results

### The gene content and organization of the PBS rod region vary between pigment types

We analysed the PBS rod region from 69 *Synechococcus* and *Cyanobium* strains (Table S1), which includes all PTs except 2B, which were extensively described elsewhere (Larsson et al. 2014; Callieri et al. 2019; Sánchez-Baracaldo et al. 2019). This dataset contains every sequenced PT 3 strain and covers a very wide range of phylogenetic diversity, with representatives of all three deep branches within *Cyanobacteria* Cluster 5 *sensu* (Herdman et al. 2001), sub-clusters (SC) 5.1 to 5.3 (Dufresne et al. 2008; Doré et al. 2020).

The PBS rod region is always situated between a phenylalanine-tRNA (tRNA-Phe_GAA_) at the 5’ end and the *ptpA* gene, which encodes a putative tyrosine phosphatase, at the 3’ end (Fig. 1 and Figs. S1-S7). Globally, the gene content and synteny of the PBS rod region *per se* (i.e., from *unk1* to *ptpA*) is remarkably conserved among cultured representatives of a given PT (Figs. S1-S7). PT 1 strains have the simplest rods and the shortest PBS rod region (the smallest is about 8 kb in *Cyanobium gracile* PCC 6307, Fig. S1). It comprises one to four copies of the *cpcBA* operon encoding α- and β-PC subunits, one PBS rod-photosystem I linker gene (*cpcL;* Watanabe et al. 2014), two to four rod linker genes (one *cpcD* and up to three copies of *cpcC*), three phycobilin lyase genes including *cpcT* (Shen et al. 2006) and the *cpcEF* operon (Fairchild et al. 1992; Swanson et al. 1992; Kronfel, Hernandez, et al. 2019), and two as-yet uncharacterized genes (*unk1-2*).

**Fig. 1:**
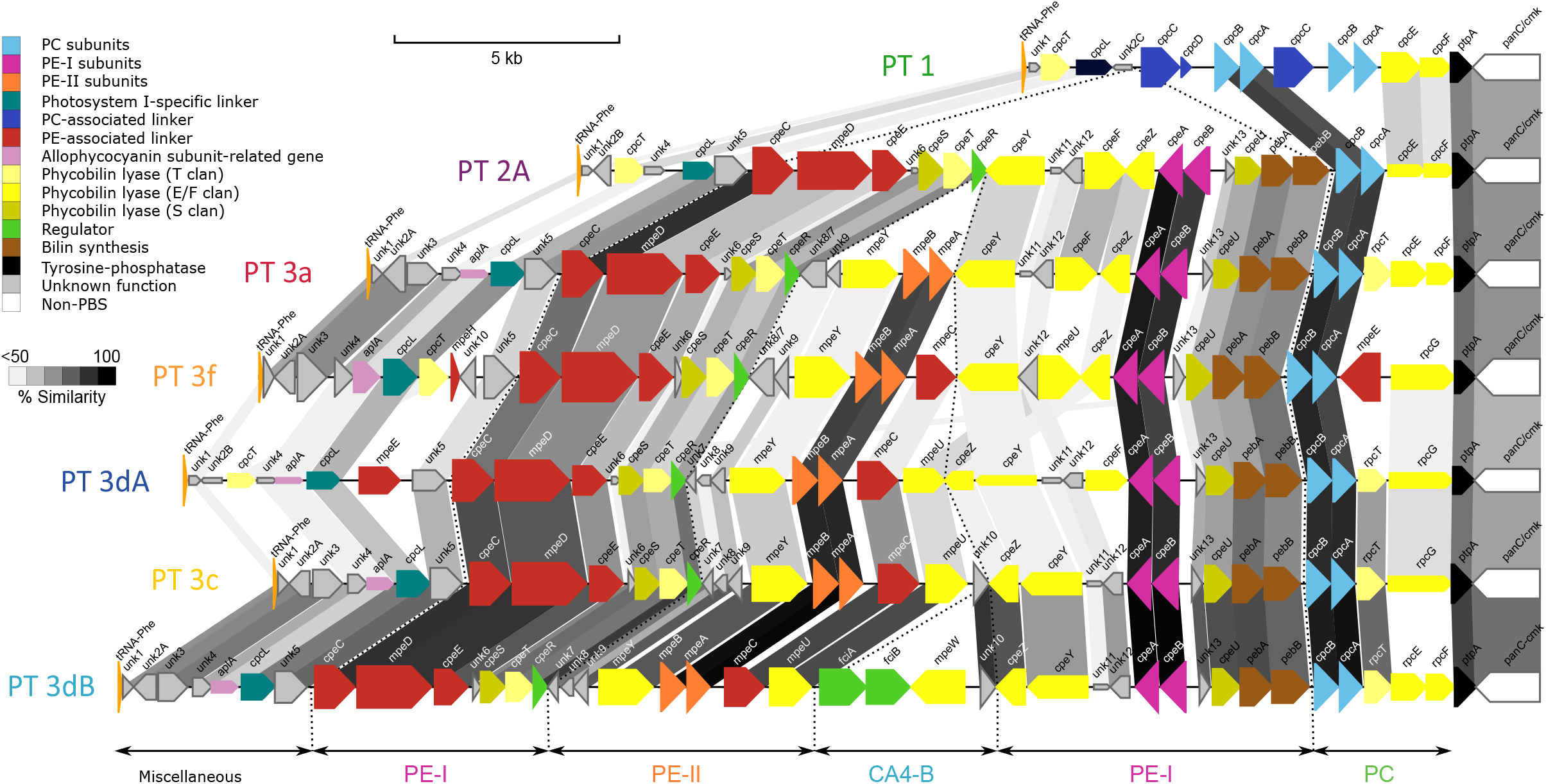
PBS rod region for strains of different pigment types. Regions are oriented from the phenylalanine tRNA (left) to the conserved low molecular weight tyrosine phosphatase *ptpA* (right). Genes are coloured according to their inferred function (as indicated in insert). Their length is proportional to the gene size and their thickness to the protein identity between strains of the same pigment type. Gray cross-links between regions are shaded according to the average protein identity between strains of two consecutive pigment types. The strains represented here are WH5701 (PT 1), WH7805 (PT 2A), RS9907 (PT 3a), KORDI-100 (PT 3f), RS9916 (PT 3dA), WH8102 (PT 3c) and A15-62 (PT 3dB). Abbreviations: PC, phycocyanin; PE-I, phycoerythrin-I; PE-II, phycoerythrin-II; PBS, phycobilisomes.

All other PTs possess a single *cpcBA* copy and no PC rod linker genes. In PT 2A, these PC genes are replaced by a set of 16 to 18 genes necessary for the synthesis and regulation of PE-I hexamers, as previously described for strain WH7805 (Six et al. 2007). This PE-specific region includes the *cpeBA* operon encoding the PE-I α- and β-subunits, three PE-associated linkers (*cpeC, cpeE* and *mpeD*), five phycobilin lyases (*cpeF, cpeS, cpeT, cpeU and cpeY;* Wilbanks and Glazer 1993b; Schluchter et al. 2010; Biswas et al. 2011; Bretaudeau et al. 2013; Kronfel, Biswas, et al. 2019; Carrigee, Mahmoud, et al. 2020), *cpeZ*, which encodes a chaperone-like protein (Kronfel, Biswas, et al. 2019), *pebA* and *pebB*, encoding the two enzymes necessary for the final two steps of PEB synthesis (Frankenberg et al. 2001), *cpeR*, encoding a PE operon activator (Cobley et al. 2002; Seib and Kehoe 2002) and a number of small, uncharacterized genes (*unk4-6* and *unk11-13*; Fig. S2).

The main difference between the PBS rod regions of PT 2A and 3a is the presence in the latter of a small cluster of five genes between *cpeR* and *cpeY*, including *unk8/7*, a fusion of *unk8* and *unk7* found as separate genes in all other PT 3 subtypes, *unk9*, another gene of unknown function, the recently characterized PEB lyase gene *mpeY* (Sanfilippo, Nguyen, et al. 2019; Grébert et al. 2021), and the *mpeBA* operon encoding the α- and β-PE-II subunits (Fig. 1 and Figs. S2-S7).

Genome composition differences between PT 3a and other PT 3 subtypes 3c, 3dA, 3dB and 3f (Mahmoud et al. 2017; Xia, Guo, et al. 2017) are mainly located in the subregion between the PE-II (*mpeBA*) and PE-I (*cpeBA*) operons (Fig. 1). All PT 3 subtypes other than 3a possess *mpeC*, encoding a PE-II associated PUB-binding linker (Six et al. 2005), inserted downstream of *mpeBA*, as well as *mpeU* encoding a partially characterized phycobilin lyase-isomerase (Mahmoud et al. 2017). Moreover, the conserved hypothetical gene *unk10*, which is absent from all PT 3a, is present in the middle of the PBS rod region of all 3c and 3dB PTs, while in PT 3dA strains it is always located in the CA4-A island, thus outside the PBS rod region. Finally, the lyase-isomerase gene *mpeQ* is present in 3c and 3dB PTs instead of the lyase gene *mpeY* (Sanfilippo, Nguyen, et al. 2019; Grébert et al. 2021).

A number of differences between PBS rod regions of various strains are more difficult to link with a specific PT. All PT 3 strains except SYN20 (PT 3a) possess the putative PE-II linker gene *mpeE*. The genomic location of this gene is highly variable, though it is generally the same among members of a given PT and clade. For example, *mpeE* can be near the 5’ or 3’ end of the PBS rod region or in a genomic location lacking genes involved in PBS structure or synthesis (Fig. S2-S7 and Table S2). Clade VII strains A15-60 and A18-25c (both PT 3c) have two slightly different copies of this linker gene, one located within and the other outside the PBS rod region. Similarly, the distribution of the putative PE-II linker genes *mpeG* and *mpeH* (the latter is a truncated version of the former, with a product length of about 59 residues for MpeH versus 312 for MpeG) cannot be linked to either a PT or a clade (Table S2). Additional examples exist for some lyase genes. For instance, all PT 3a strains contain the *rpcEF* operon, encoding the two subunits of a C84 α-PC PEB lyase, upstream of *ptpA* (Swanson et al. 1992; Zhou et al. 1992). Yet other PT 3 subtypes may have either the *rpcEF* operon or *rpcG*, a fusion gene that encodes a C84 α-PC PEB lyase-isomerase and was thought to confer *Synechococcus* cells a better fitness to blue light environments (Blot et al. 2009). This interchangeability is found even between closely related strains of the same PT and clade, such as for the clade CRD1/PT 3dA strains GEYO and MIT S9220 or the clade II/3c strains CC9605 and RS9902 (Fig. S5) and strain MINOS11 (SC 5.3/PT 3dB) even possesses both genes (Fig. S7) Similarly, RS9916 possesses both *cpcT* and *rpcT*, two enzymes that potentially act on the same site (C153 β-PC) but attach PCB and PEB chromophores, respectively (Shen et al. 2006; Blot et al. 2009). Some additional variations of ‘typical’ PBS rod regions are also worth noting. Out of five PT 2A strains, only CB0205 and A15-44 possess the allophycocyanin-like gene *aplA* (Montgomery et al. 2004) which, as in all PT 3 strains, is located between *unk4* and *cpcG2* (Fig. S2 and Table S2). CB0205 also has a unique insertion of nine genes of unknown function, including *unk3*, in the middle of its PBS rod region between *unk12* and *cpeF*, as observed in some natural PT 2A populations from the Baltic Sea (Larsson et al. 2014).

Finally, a number of genomes were found to possess a DNA insertion of variable size between the tRNA-Phe_GAA_ and *unk1* (Figs. S1, S3, S5-S7). The most extreme example of this is in TAK9802, where the 22.8 kb DNA insertion is almost as large as the 24.5 kb PBS rod region. These insertions have striking similarities to ‘tycheposons’, a novel type of mobile genetic elements that have been found to be responsible for the translocation of 2-10 kbp fragments of heterologous DNA in *Prochlorococcus* (Hackl et al. 2020). Indeed, the hallmarks of tycheposons include the systematic presence of a tRNA at the 5’ end of the insertion, the localization of this insertion upstream a genomic island important for niche adaptation —in the present case, adaptation to light color—, and the presence in the DNA insertion of a variety of mobile elements. These notably include a putative tyrosine recombinase (here, the purple-circle gene in Figs. S1, S3, S6 and S7, which corresponds to Cyanorak CLOG CK_00022444; Garczarek et al. 2021).

While the genomic organization of the PBS rod region *per se*, i.e. excluding the tycheposon, is broadly conserved, we found a high level of allelic diversity of the genes of this region and the proteins they encode (Figs. 1 and S8). The most conserved are genes encoding the α- and β-subunits of phycobiliproteins. The sequence of each of these proteins is at most only about 10% different from its closest ortholog. The sequences of the linker proteins show greater variation between strains, with some having less than 70% identity to their closest ortholog. This variability is even greater for phycobilin lyases and uncharacterized conserved proteins, with some showing less than 60% sequence identity to their closest orthologs (Fig. S8). Such highly divergent sequences may in some cases reflect functional differences, as was recently demonstrated for MpeY and MpeQ (Grébert et al. 2021) and for CpeF and MpeV (Carrigee, Frick, et al. 2020).

### Targeted metagenomics unveils PBS rod regions from field populations

We investigated the genetic variability of PBS rod regions from natural *Synechococcus* populations using a targeted metagenomic approach that combined flow cytometry, cell-sorting, WGA and fosmid library screening (Humily et al. 2014). This method enabled us to retrieve PBS rod regions from natural populations in the North Sea, northeastern Atlantic Ocean and various locations within the Mediterranean Sea (Fig. 2A and Table S3). In addition, the high-resolution phylogenetic marker *petB* (Mazard et al. 2012) was sequenced to examine the phylogenetic diversity of these natural *Synechococcus* populations. Samples collected from the North Sea (fosmid libraries H1-3 in Fig. 2B) and English Channel (library A) were exclusively composed of the cold-adapted clade I, mostly sub-clade Ib. Samples from the northeastern Atlantic Ocean were co-dominated by CRD1 and either clade I (libraries G1 and G2) or the environmental clades EnvA and EnvB (library E; EnvB is sometimes called CRD2; (Ahlgren et al. 2019). *Synechococcus* populations from the western Mediterranean Sea were largely dominated by clade III (exclusively of sub-clade IIIa) at the coastal ‘Point B’ station located at the entrance of the Bay of Villefranche-sur-Mer (https://www.somlit.fr/villefranche/; library F), while they essentially consisted of clade I (mostly sub-clade Ib) at station A of the BOUM cruise (library I2; Moutin et al. 2012) and at the long-term monitoring station BOUSSOLE located in the Gulf of Lions (library I1; Antoine et al. 2008). Eastern Mediterranean Sea populations collected at BOUM stations B and C were mainly from clades III, with sub-clade IIIa dominating. These large differences in clade composition reflect the distinct trophic regimes of the sampled sites and the diversity patterns observed here are globally consistent with previous descriptions of the biogeography of *Synechococcus* clades (Zwirglmaier et al. 2008; Mella-Flores et al. 2011; Paulsen et al. 2016). In particular, CRD1 and EnvB are known to co-occur in iron-poor areas (Farrant et al. 2016; Sohm et al. 2016) and the northeastern Atlantic Ocean has been reported to be iron-limited (Moore et al. 2013).

**Fig. 2:**
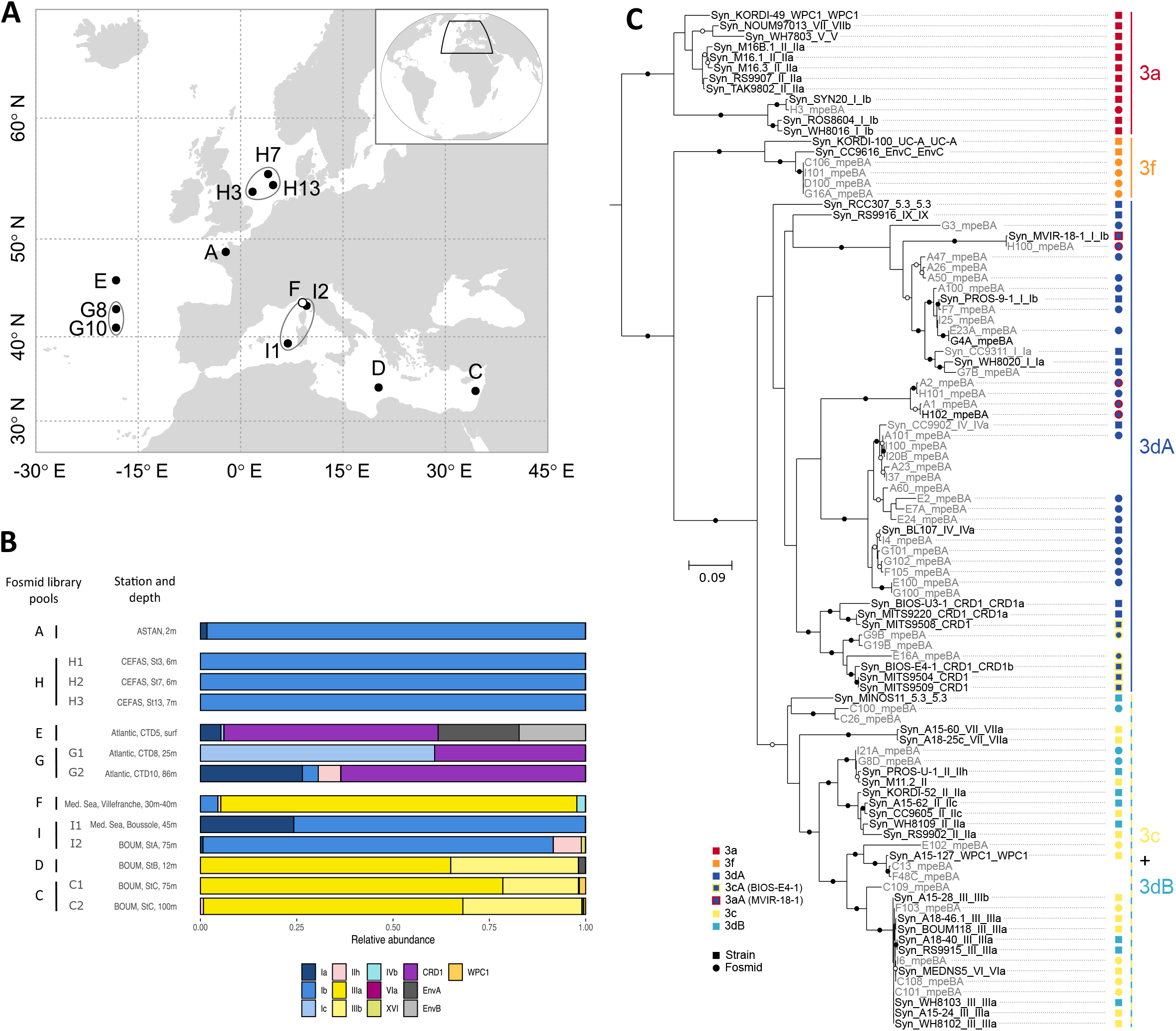
Characterization of PBS rod regions from natural population of *Synechococcus*. (*A)* location of sampling sites used for fosmid library construction. (*B*) *Synechococcus* genetic diversity at each station, as assessed with the phylogenetic marker *petB*. (*C*) *mpeBA* phylogeny for isolates (black) and fosmids (gray). Squares and circles on right hand side correspond to reference strains and fosmids, respectively.

To obtain the largest possible diversity of PBS rod regions from field *Synechococcus* populations, fosmid libraries generated from similar geographic areas and/or cruises and showing comparable relative clade abundance profiles were pooled (as indicated in Figs. 2A-B and Table S3) before screening and sequencing. Assembly of the eight fosmid library pools resulted in the assembly of 230 contigs encompassing either a portion or all of the PBS rod region. These contigs were an average size of approximately 5.5 kb and each library produced at least one contig longer than 10 kb (Table S4). Each contig was assigned to a PT based on its genomic organization and similarity to PBS rod regions of characterized strains (Fig. 1 and Figs. S1-7). Most contigs corresponded to PT 3dA (145 out of 230), but all PT 3 subtypes represented in our reference dataset were found. There were five PT 3a contigs, 18 PT 3f contigs and 62 PT 3c/3dB contigs, of which nine could be unambiguously attributed to PT 3c and five to PT 3dB, based on the absence or presence of a CA4-B genomic island between *mpeU* and *unk10*, respectively (Fig. 1).

A representative selection of the most complete contigs is provided in Figure 3. Most contigs are syntenic with PBS rod regions from characterized strains, notably those assigned to PT 3a retrieved from the North Sea (contig H3), or those assigned to PT 3f either retrieved from the northeastern Atlantic Ocean (contig G16A) or from the Mediterranean Sea (all other contigs assigned to PT 3f; Fig. 3B). Yet, a number of contigs assigned to PT 3dA (E101, F100 and F101; Fig. 3A) exhibit a novel gene organization compared to reference 3dA strains characterized by the insertion of a complete CA4-A genomic island between *unk3* and *unk4* at the 5’-end of the PBS rod region (Fig. S5). Five other contigs (G100, E28, E20, F12 and H104) apparently have the same organization since they possess a complete or partial *mpeZ* gene located immediately upstream of *unk4*. Despite this novel location, the genes of this CA4-A island are phylogenetically close to those of the PT 3dA/clade IV strains CC9902 or BL107 (Fig. 3A). This new arrangement is found in contigs from different sequencing libraries and geographically diverse samples, so it is most likely not artefactual. Contigs corresponding to the canonical 3dA PBS rod region (Fig. 1) were also found in most libraries and contain alleles most similar to those of PT 3dA strains from either clade I or IV (Fig. 3B).

**Fig. 3:**
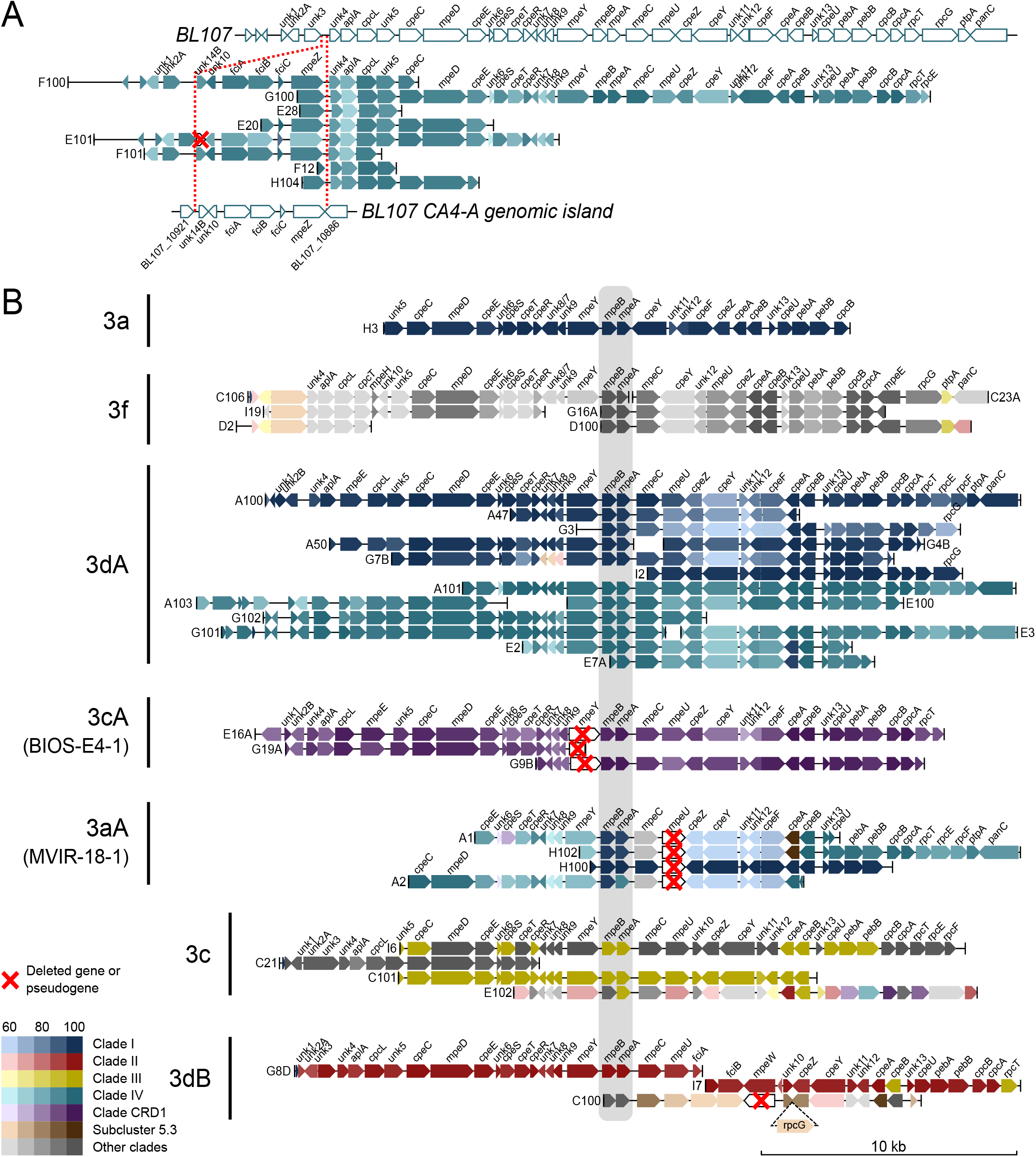
partial or complete PBS rod region retrieved from natural populations. (*A*) Description of a new genomic organization related to 3dA pigment type with the CA4-A genomic island inserted at the 5’-end of the PBS rod genomic region. The PBS rod and CA4-A genomic regions of strain BL107 (3dA/clade IV) is shown as a reference. (*B*) Contigs other than those in (*A*) and longer than 10 kb, sorted according to their organization and inferred corresponding pigment type. Colors represent the clade of the strain giving the best BlastX hit within the given pigment type. The highly conserved *mpeBA* operon is shaded in gray.

Several contigs from the English Channel and North Sea (A1, A2, H100 and H102) that displayed a similar gene organization to PT 3a strains lacked *mpeU* (Fig. 3B), which encodes a lyase-isomerase that attaches PUB to an as-yet undetermined residue of PEII (Mahmoud et al. 2017). The absence of *mpeU* was also observed in the genome of strain MVIR-18-1 (clade I/PT 3a; Fig. S5A), which was shown to display a constitutively low PUB:PEB ratio (Humily et al. 2013). MVIR-18-1 was isolated from the North Sea, suggesting that natural populations representative of this *mpeU*-lacking variant may be common in this area. Similarly, the gene organization of three contigs originating from the northeastern Atlantic Ocean and assigned to CRD1 (E16A, G19A and G9B; Fig. 3B) matched an unusual PBS rod region found in five out of eight reference CRD1/PT 3dA strains (Fig. S5A). In these strains, the *mpeY* sequence is either incomplete (in MITS9508) or highly degenerate (in BIOS-E4-1, MITS9504, MITS9509 and UW179A) and *fciA* and *fciB*, which encode CA4 regulators, are missing (Fig. S5B), resulting in a PT 3c phenotype (Humily et al. 2013). This particular genomic organization was recently suggested to predominate in CRD1 populations from warm high nutrient-low chlorophyll areas, in particular in the South Pacific Ocean (Grébert et al. 2018). Thus, these contigs provide additional, more compelling evidence of the occurrence of these natural variants in field populations of CRD1.

A number of contigs corresponding to the phylogenetically indistinguishable PBS rod regions of the 3c and 3dB PTs were assembled from samples from the Mediterranean Sea and the northeastern Atlantic Ocean (Fig. 2B). Among these, five (C100, C40, G8D, I7 and I21) could be assigned to PT 3dB due to the presence of a CA4-B genomic island between *mpeU* and *unk10* (Fig. 1). The gene organization of contig C100, whose alleles are most similar to the PT 3dB/SC 5.3 strain MINOS11, closely resembles the unique PBS rod region of this strain, which has an additional *rpcG* gene between *unk10* and *cpeZ* (Fig. S7). Interestingly, the CA4-B genomic island of C100 lacks *mpeW* (Fig. 2B) and thus may represent a novel natural variant. In contrast with the genes of contigs assigned to PT 3dA, which are closely similar to clades I, IV or CRD1, those assigned to PT 3c and 3dB are most closely related to representatives of different clades, including clades II, III, WPC1 and 5.3. These distinct clade/PT combinations corroborate previous observations made in culture and in the field (Grébert et al. 2018) and are consistent with the population composition observed with *petB* at the sampling locations of these libraries (Fig. 2B). Of note, some contigs from fosmid library E (e.g., contig E102; PT 3c in Fig. 3B) possessed alleles that were highly divergent from all reference strains and likely belong to the uncultured clades EnvA or EnvB, which together represented more than 30% of *Synechococcus* population in sample E (Fig. 2B).

We augmented these field observations by examining the PBS rod regions contained within the publicly available single-cell amplified genomes (SAGs) of marine *Synechococcus* (Berube et al. 2018). Out of 50 *Synechococcus* SAGs, eleven corresponded to PT 3c, three to PT 3dB, two to either PT 3c or 3dB, and 17 to PT 3dA (Fig. S9 and Table S1). Using a core genome phylogeny based on a set of 73 highly conserved markers (Table S5 and Fig. S10), we determined that these SAGs belong to clades I, II, III, IV, CRD1 as well as some rare clades, including one to EnvA, one to XV, three to SC5.3, and seven to EnvB. All of the EnvB SAGs contained a PBS rod region which was very similar to PT 3c except for the insertion of *rpcG* between *unk10* and *cpeZ*. Due to the absence of any sequenced EnvB isolate in our reference database, genes from these regions appear most similar to genes from a variety of strains and clades. The same is true for SAGs from clades EnvA (AG-676-E04) and XV (AG-670-F04). All SAGs assigned to PT 3dB belong to SC5.3 and possess a PBS rod region similar to that of MINOS11 except for a ca. 15 kb insertion between the tRNA-Phe_GAA_ and *unk2* in AG-450-M17 (Fig. S9B). The ten SAGs within clade CRD1 all belong to PT 3dA, but only one of these contains an intact *mpeY* gene (Fig. S9C). The three clade I SAGs also belong to PT 3dA. Of these, AG-679-C18 provides the first example of a *mpeY* deficiency outside of clade CRD1, further highlighting the prevalence of BIOS-E4-1-like populations, which are phenotypically similar to (but genetically distinct from) PT 3c, in the environment (Grébert et al. 2018). Interestingly, all four clade IV SAGs contain the novel PT 3dA arrangement found in several fosmids, where the CA4-A genomic island is located in the PBS rod region between *unk3* and *unk4*. Finally, a number of the SAGs have additional genes between the tRNA-Phe_GAA_ gene and *unk1*, including recombinases, restriction enzymes, etc., sometimes in multiple copies, for example in the clade II SAGs AG-670-A04 and AG-670-B23 (Fig. S9A).

Altogether, the PBS rod regions retrieved from both fosmids and SAGs were very diverse and some contained alleles whose sequences were highly diverged from those of sequenced isolates. As was found in our comparative analysis of strains, the sequence diversity for lyases and linker proteins was much higher than for phycobiliproteins (Fig. S8).

### Genes within the PBS rod region have a very different evolutionary history than the rest of the genome

Several phylogenetic analyses based on phycobiliprotein coding genes have shown that strains tend to group together according to their PT rather than their vertical (core) phylogeny. The *cpcBA* operon enables discrimination between PT 1, 2A, 2B and 3, while the *cpeBA* operon allows separation of PT 2A, 3a, 3dA, 3f and the 3c+3dB group and the *mpeBA* operon is best for distinguishing between the PT 3 subtypes (Everroad and Wood 2012; Humily et al. 2014; Xia, Partensky, et al. 2017; Grébert et al. 2018; Xia et al. 2018). By creating a *mpeBA* phylogenetic tree using all available genomes from *Synechococcus* PT 3 strains, we confirmed that alleles within a given PT 3 subtype are more closely related to one other than they are to other PTs from the same clade (Fig. 4, left tree). However, we also observed that within each *mpeBA* clade, the tree topology actually resembles the topology based on the vertically transmitted core gene *petB* (Fig. 4, right tree). The few exceptions to this finding could correspond to inter-clade horizontal gene transfers. The most striking example of this is the clade VI strain MEDNS5, which seemingly possesses a clade III PT 3c/3dB-like *mpeBA* allele (Fig. 4).

**Fig. 4:**
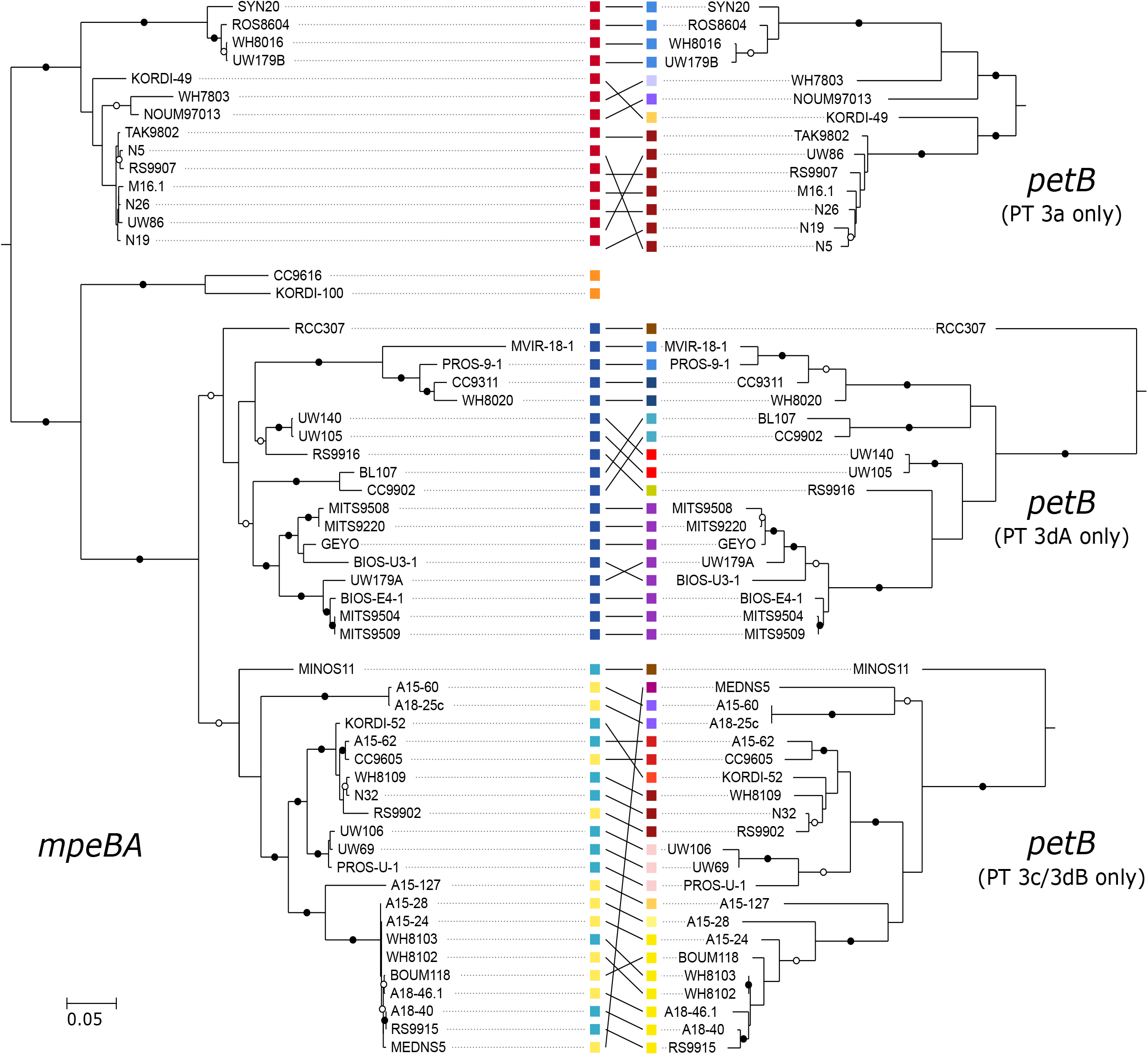
Correspondence between phylogenies for the *mpeBA* operon and the marker gene *petB*, which reproduces the core genome phylogeny. The pigment type for each strain is indicated by a coloured square in the *mpeBA* phylogeny, and its clade similarly indicated in the *petB* phylogeny.

In order to explore the evolution of the PBS genes in greater detail, we used ALE (Szöllősi et al. 2013) to reconcile phylogenetic trees for each gene present in the core genome with the species tree inferred from a set of 73 core genes (Table S5 and Fig. S10). This comparison allows the inference of evolutionary events such as duplications, horizontal transfers, losses and speciations that can best explain the observed gene trees in light of the evolution of the species (Fig. 5). Genes from the PBS rod region experienced significantly more transfers (on average 23.5 vs. 14.8 events per gene; Wilcoxon rank sum test p=1.2×10^−13^) and losses (38.5 vs. 27.5; p=1.7×10^−11^) than other genes in the genome, with no significant difference in gene duplications and speciations (Fig. 5 and Table 1). Consistent with the observation that genes within the PBS rod region are single copy except for *cpcABC* in PT 1, the increase in transfers was very similar to the increase in losses. We conclude that such transfers actually correspond to allelic exchange, whereby homologous recombination mediates the replacement of one allele by another.

**Fig. 5:**
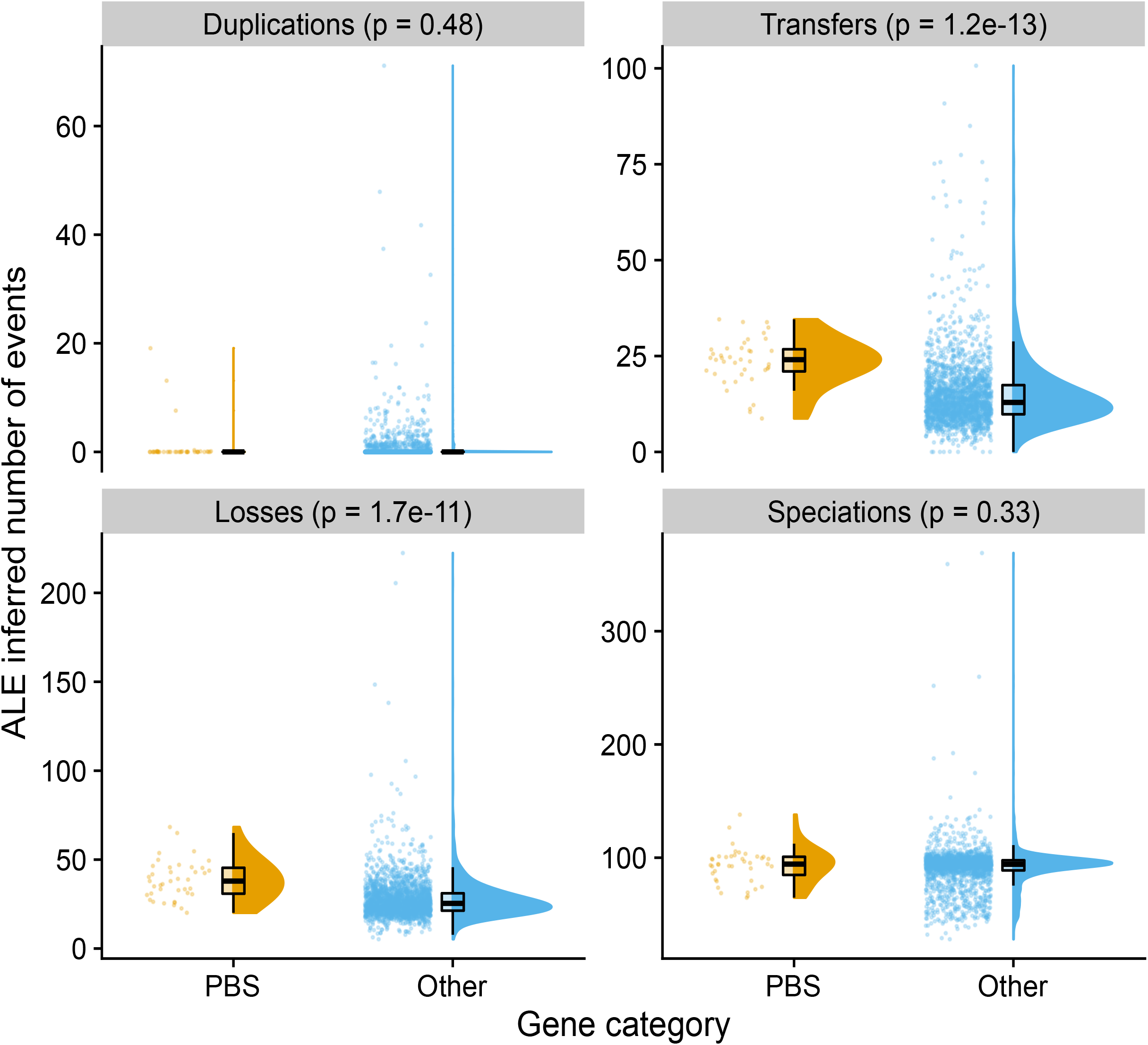
Evolutionary events affecting genes present in more than half of the analysed genomes inferred by reconciling gene trees with the species tree. Genes were classified either as belonging to the PBS rod region (“PBS genes”) or as other genes (“Other”). *P*-values for Wilcoxon rank sum exact test are shown.

**Table 1:** Frequency of transfer events inferred by ALE for genes of the PBS rod region.

| Gene category | Evolutionary event | Min. | Median | Mean | Max. |
| --- | --- | --- | --- | --- | --- |
| PBS | Duplications | 0.0000 | 0.0000 | 1.0190 | 19.0900 |
| PBS | Transfers | 8.72 | 24.05 | 23.52 | 34.57 |
| PBS | Losses | 20.10 | 37.92 | 38.50 | 68.31 |
| PBS | Speciations | 64.70 | 94.36 | 93.08 | 138.01 |
| Other | Duplications | 0.000 | 0.000 | 0.555 | 71.110 |
| Other | Transfers | 0.000 | 12.910 | 14.801 | 100.680 |
| Other | Losses | 5.22 | 25.38 | 27.53 | 222.40 |
| Other | Speciations | 28.01 | 94.19 | 91.32 | 369.02 |

Finer-grained analysis of transfer events showed that they are slightly more frequent within clades for PBS genes than for other genes (9.7 vs. 8.4, p=1.1×10^−3^; Table 2 and Fig. S11), and more than twice as frequent between clades (13.9 vs. 6.1, p=10^−15^; Table 2 and Fig. S11). As a result, most transfer events identified for PBS genes occurred between clades, whereas other genes were primarily transferred within the same clade (Fig. S11). We then analyzed transfer events inferred for genes within the PBS rod region by their direction (Table S6). The most frequent recipient of transfer (frequency of 29.07) was strain N32, which can be explained by the fact that this PT 3dB strain is within a lineage of clade II that is otherwise solely made up of PT 3a strains (node 177; Fig. S10). The second most frequent transfer recipient was strain MEDNS5 (clade VI; frequency=23.71), which possesses the allele of a PT 3c/3dB strain of clade III, as noted previously (Fig. 4). Strains RS9902 and A15-44, both within clade II, were the third (21.69) and fourth (18.79) most frequent recipient strains. A15-44 belongs to PT 2, a quite rare PT among strictly marine SC 5.1 *Synechococcus* strains (Grébert et al. 2018), and indeed groups in the tree with the PT 3c strain RS9902 (Fig. S10). The apparent high transfer frequency to RS9902 could thus represent transfers of PT 2 to the ancestor of RS9902 and A15-44 (node 103 in Fig. S10 and Table S6), followed by transfers of PT 3c to RS9902. Other strains for which high frequency of transfers were inferred are KORDI-49 (WPC1/3aA), RCC307 (SC 5.3/3eA), WH7803 (V/3a), CB0205 (SC 5.2/2), which all represent rare combinations of clade and PT (Fig. S10 and Table S6). Internal nodes with high frequency of transfers were mostly deep-branching, representing the ancestor of SC 5.3, clades I, V, VI, VII and CRD1, 5.1B, and 5.1A (nodes 171, 189, 191, 197, with frequencies of 22.2, 14.5, 13.9 and 12.9, respectively). Taken together, these results indicate that genes of the PBS rod region have a very ancient evolutionary history marked by frequent recombination events both within and between *Synechococcus* clades.

**Table 2:** Frequency of transfer events inferred by ALE for genes of the PBS rod region.

| Gene category | Transfer event | Min. | Median | Mean | Max. |
| --- | --- | --- | --- | --- | --- |
| PBS | Intra-clade | 5.260 | 9.655 | 9.655 | 13.410 |
| PBS | Extra-clade | 2.08 | 14.57 | 13.86 | 21.29 |
| PBS | Intra-PT | 3.910 | 11.685 | 11.405 | 19.770 |
| PBS | Extra-PT | 4.51 | 12.74 | 12.11 | 17.42 |
| Other | Intra-clade | 0.000 | 8.305 | 8.419 | 40.070 |
| Other | Extra-clade | 0.000 | 4.325 | 6.143 | 81.730 |
| Other | Intra-PT | 0.000 | 5.970 | 6.292 | 40.570 |
| Other | Extra-PT | 0.000 | 6.935 | 8.269 | 68.060 |

## Discussion

### The variable gene content of the PBS rod region relies on a tycheposon-like mechanism

Most of the current knowledge about the genomic organization and genetic diversity of the *Synechococcus* PBS rod region has relied on analysis of the first 11 sequenced genomes (Six et al. 2007) and later analyses of metagenomic assemblages or strains retrieved from a few specific locations. These include the SOMLIT-Astan station in the English Channel (Humily et al. 2014), the Baltic and Black Seas, where a new organization of the PBS rod region (PT 2B) was found associated with SC 5.2 populations (Larsson et al. 2014; Callieri et al. 2019), and freshwater reservoirs dominated by PT 2A/SC 5.3 populations (Cabello-Yeves et al. 2017). Here, we have analyzed a wide set of *Synechococcus* and *Cyanobium* genomes (69 genomes and 33 SAGs; Table S1) as well as PBS rod regions directly retrieved from a variety of trophic and light environments (229 contigs; Table S4). Together, these cover all of the genetic and PT diversity currently known for this group (except PT 2B), enabling us to much better assess the extent of the diversity within each PBS rod region type.

While our data confirmed that the gene content and organization of this region are highly conserved within a given PT independent of its phylogenetic position (Six et al. 2007), they also highlighted some significant variability within each PT. We notably unveiled a novel and evolutionary important trait of PBS rod regions, namely the frequent presence of DNA insertions between the tRNA-Phe_GAA_ and *unk1* in both strains or SAGs (Figs. S1-S7 and S9). While these insertions share striking similarities to ‘tycheposons’, a novel type of mobile genetic elements recently discovered in *Prochlorococcus* and notably the presence in many of them of a complete or partial tyrosine recombinase (Hackl et al. 2020), they also display some specificities. Indeed, *Prochlorococcus* tycheposons have been associated with seven possible tRNA types (Hackl et al. 2020), but never with a tRNA-Phe_GAA_, which is the tRNA type systematically found upstream of PBS rod regions. Also, while in *Prochlorococcus* the tyrosine recombinase is most often located immediately downstream the tRNA at the 5’ end of the tycheposon, in *Synechococcus* it is found at the distal end of the tycheposon. Closer examination of this distal region in several *Synechococcus* genomes (notably in TAK9802) revealed a remnant of tRNA-Phe_GAA_ located between the tyrosine recombinase gene and *unk1* in genomes where the former gene was complete. This observation strongly suggests that insertion of DNA material occurred by homologous recombination via a site-specific integrase at the level of this tRNA, since this process often leaves the integrated elements flanked on both sides with the attachment site motif (Grindley et al. 2006). The presence of other recombinases in the DNA insertion in a few *Synechococcus* strains (e.g., a putative site-specific, gamma-delta resolvase in TAK9802, N5 and M16.1; red-contoured gene in Fig. S3A) may have resulted from multiple integrations at the same tRNA site, a phenomenon also reported for *Prochlorococcus* tycheposons (Hackl et al. 2020). Altogether, these observations strongly suggest that the wide genomic diversity of PBS rod regions could involve a tycheposon-like mechanism, promoting the import of foreign DNA as flanking material to mobile genetic elements, some of this material having been retained by natural selection and used for adaptation to specific light color niches.

### Novel insights into the evolution of CA4 islands and chromatic acclimation

The characterization of many PBS rod regions from the environment that were retrieved from SAGs or fosmids led us to the discovery of a new location for the CA4-A genomic island near the 5’-end of the PBS rod region (Fig. 3A and S9C) rather than elsewhere, as found in the genomes of all 3dA strains sequenced thus far (Fig. S5B). All of the genes of the PBS rod region containing these atypical CA4-A islands have strongest similarity to the corresponding genes in PT 3dA strains. Therefore, this new organization is unlikely to correspond to a new PT phenotype/genotype but rather to a previously unidentified PT 3dA variant. The localization of the CA4-A region at the 5’-end of the PBS rod region supports the abovementioned hypothesis that the increases in the complexity of *Synechococcus* pigmentation by progressive extension of the PBS rod region primarily occurred via a tycheposon mechanism. This finding also provides a simple solution to the paradox of how two physically separate genomic regions encoding related components, with evolutionary histories that differ from the rest of the genome, were still able to co-evolve.

The CA4-A island not only has a highly variable position within the genome, but its gene content is also variable. Indeed, five out of eight CRD1 strains with a 3dA configuration of the PBS rod region possess an incomplete CA4-A island (Fig. S5B). In contrast, the CA4-B island is always found at the same position in the genome and is complete in all 3dB strains sequenced so far (Fig. S7). If the CA4-B island also has been generated in a tycheposon, the mechanism by which it has been transposed in the middle of the PBS rod region remains unknown, since there are no known recombination hotspots in this region. Contig C100, which appears to lack the PEB lyase-encoding gene *mpeW* (Fig. 3B; Grébert et al. 2021), is the first documented example of an incomplete CA4-B region. We predict that this new genotype has a constitutively high PUB:PEB ratio since this organism is likely to contain an active lyase-isomerase MpeQ (Grébert et al. 2021). It also has a *rpcG* gene encoding a PC lyase-isomerase (Blot et al. 2009) located immediately downstream of the CA4-B island and on the same strand as *unk10* (Figs. 3B and S7, 3dB). This arrangement, which has thus far only been observed in MINOS11, suggests that *rpcG* expression could be controlled by light color and its protein product compete with those encoded by the *rpcE-F* operon, since both act on the same cysteine residue of α-PC (Swanson et al. 1992; Zhou et al. 1992; Blot et al. 2009). This arrangement would be similar to the relationship between *mpeZ* and *mpeY* (Sanfilippo, Nguyen, et al. 2019) or between *mpeW* and *mpeQ* (Grébert et al. 2021). If confirmed, this would be the first case of chromatic acclimation altering the chromophorylation of PC instead of PE-I and PE-II in marine *Synechococcus*.

### A hypothesis for the evolution of the PT 3 PBS rod region

The apparent mismatch between PBS pigmentation and vertical phylogeny raises the intriguing question of how different PTs have evolved and been maintained independently from an extensive clade diversification. It is generally agreed that the occurrence of the *mpeBA* operon in marine *Synechococcus* spp. PT 3 and the closely related uncultured *S. spongiarum* group resulted from gene duplication and divergence of a pre-existing *cpeBA* operon (Apt et al. 1995; Everroad and Wood 2012; Sánchez-Baracaldo et al. 2019). Yet phycobiliproteins are part of a complex supramolecular structure, interacting with many other proteins such as linkers, phycobilin lyases and regulators, which all need to co-evolve. Here, we propose an evolutionary scenario of progressively increasing complexity for the diversification of PT 3 from a PT 2/3 precursor (Fig. 6). Our model integrates recent advances in our understanding of the functional characterization of PBS gene products, notably phycobilin lyases (Shukla et al. 2012; Sanfilippo et al. 2016; Mahmoud et al. 2017; Kronfel, Biswas, et al. 2019; Sanfilippo, Nguyen, et al. 2019; Carrigee, Frick, et al. 2020; Grébert et al. 2021). Our proposal does not include PT 2B since strains exhibiting this PT generally possess several phycocyanin operons, such as PT 1 (Callieri et al. 2019), and it cannot be established with certainty whether of PT 2B or PT 2A occurred first.

**Fig. 6:**
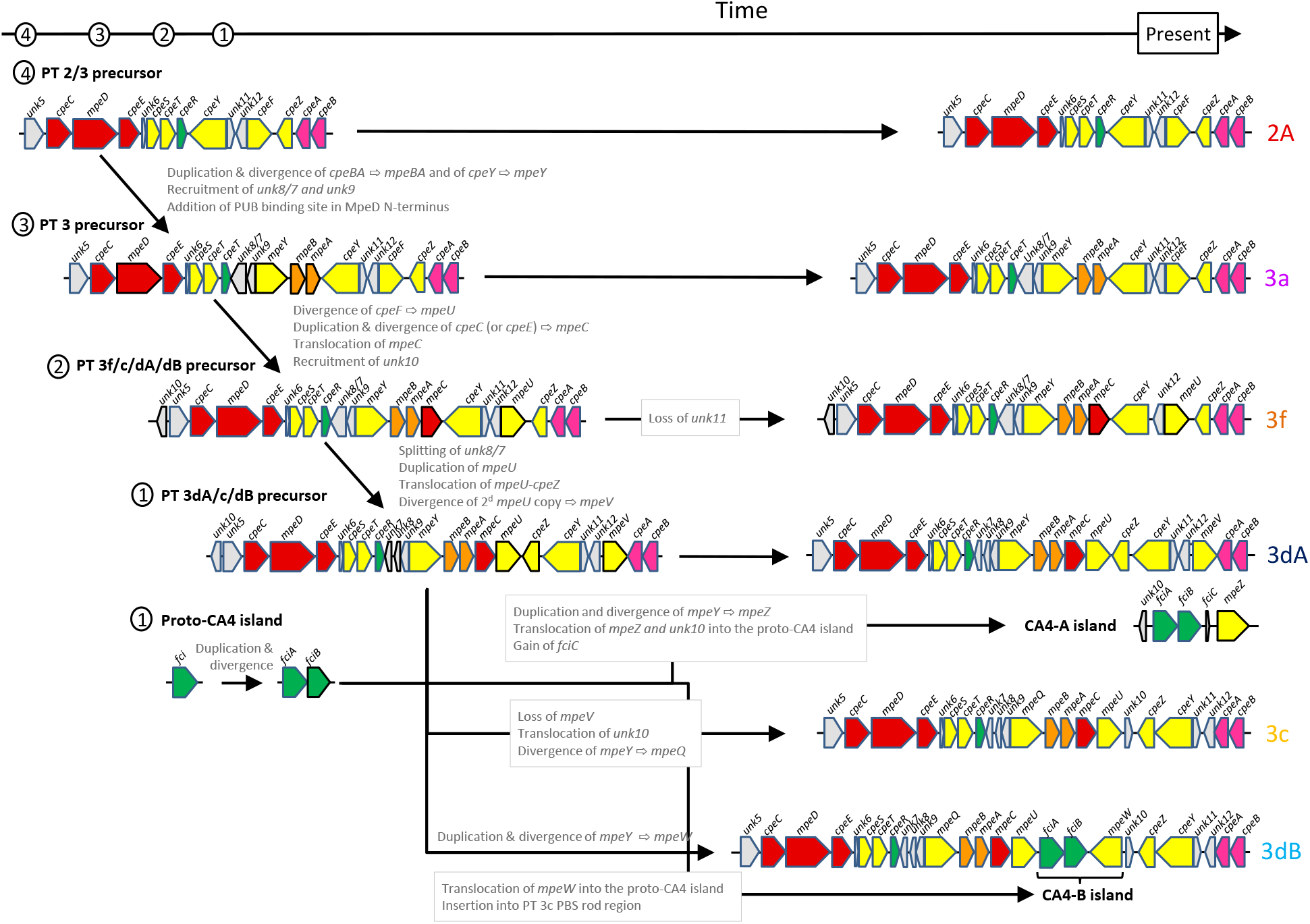
Putative evolutionary scenario for the occurrence of the different *Synechococcus* PT 3 subtypes. This scenario is congruent with individual phylogenies of genes in the PBS rod region. Note that the 5’- and 3’-end of the PBS rod region are cropped for better visualisation of the PE-I/PE-II sub-regions. Genes that changed between two consecutive PT precursor steps are highlighted by black contours (instead of blue for the other genes).

The first step toward PE-II acquisition by the PT 2/3-like precursor involved the generation of a *mpeBA* operon precursor by duplication and divergence from an ancestral *cpeBA* operon. This was accompanied by the concomitant duplication of the ancestral the PE-I specific lyase gene *cpeY* and its divergence to a precursor of the PEII-specific *mpeY* lyase gene (Fig. 6). The origin of the *unk8/7* fusion gene and *unk9*, occurring at the 5’-end of the PE-II subregion in all PT 3 strains (Fig. 1 and Fig. S2), is more difficult to assess. However, it is noteworthy that Unk9 and the two moieties of Unk8*/7* all belong to the Nif11-related peptide (N11P) family, which shows extensive paralogous expansion in a variety of cyanobacteria (Haft et al. 2010). Although some members of the N11P family have been suggested to be secondary metabolite precursors (Haft et al. 2010; Tang and van der Donk 2012; Cubillos-Ruiz et al. 2017), the functions of the Unk8/7 and Unk9 peptides remains unclear. Yet the localization of their genes in the PE-II sub-region of all PT 3 strains strongly suggests a critical role in PE-II biosynthesis or regulation. One possibility is that they modulate the specificity of some PE-I lyases to extend their activity to PE-II subunits. Another, more subtle, change that occurred during the evolution of PT 3 was the change in the N-terminal part of the MpeD linker to include a specific insertion of 17 amino acids near the N-terminal region that is involved in PUB binding, as found in all PEII-specific linkers (Six et al. 2005). Present-day PT 3a would have directly descended from this PT 2/3 last common ancestor (LCA). Accordingly, the PBS rod region from PT 3a is the simplest of all PT 3 and PT 3a sequences form the most basal clade in both *mpeBA* and *mpeWQYZ* phylogenies when these are rooted using *cpeBA* and *cpeY* sequences respectively (Fig. 2C).

The three main differences between the PT 2/3 LCA and other PT 3 LCAs are the acquisition of the linker gene *mpeC*, which most likely resulted from the duplication and divergence of a pre-existing PE-I linker (either *cpeC* or *cpeE*), the acquisition of *unk10*, encoding an additional member of the N11P family, and the replacement of *cpeF* by *mpeU* (Fig. 6). The lyase-isomerase MpeU belongs to the same family as the PEB lyase CpeF and is likely to have been derived from it. Even though the CpeF/MpeU phylogeny is unclear due to the deep tree branches having low bootstrap supports (Mahmoud et al. 2017; Carrigee, Frick, et al. 2020), we hypothesize that the PT 3f/c/dB/dA precursor already had a *mpeU*-like gene. Phylogenetic trees of *mpeBA* and of the *mpeWQYZ* enzyme family places the recently described PT 3f (Xia et al. 2018) in a branch between those formed by PT 3a in one case and PT 3dA and 3c/3dB sequences in the other (Fig. 2C). Consistently, the organization of the PT 3f PBS rod region appears to be intermediate between PT 3a and the more complex PT 3c/3dB/3dA regions. The only difference between the PT 3f/c/dB/dA precursor and the present-day PT 3f would be the loss of *unk11*, a short and highly variable open reading frame (Fig. 6).

The PT 3f/c/dB/dA LCA then would have evolved to give the common precursor of PT 3dA/c/dB. This step likely involved four events: i) the splitting of the *unk8/7* gene into two distinct genes, *unk8* and *unk7*, ii) the duplication of *mpeU* followed by iii) a tandem translocation of one *mpeU* gene copy and *cpeZ* between *mpeC* and *cpeY*, and iv) the divergence of the second *mpeU* copy to give *mpeV*, encoding another recently characterized lyase-isomerase of the CpeF family (Carrigee, Frick, et al. 2020). Again, the poor bootstrap support of deep branches of the CpeF/MpeU/MpeV phylogeny (Carrigee, Frick, et al. 2020) makes it difficult to confirm this hypothesis, and we cannot exclude the possibility that *mpeV* was derived directly from *cpeF*. Since the 3dA-type PBS rod region does not exist in present-day *Synechococcus* spp. without co-occurrence of a CA4-A island, the proto-CA4 island must also have evolved concurrently with the PT 3dA precursor. The two regulatory genes it contains, *fciA* and *fciB*, likely originate from the duplication and divergence of an ancestral *fci* precursor gene encoding a member of the AraC family. Both FciA and FciB possess a AraC-type C-terminal helix-turn-helix domain, yet their N-terminal domains have no similarity to any known protein (Humily et al. 2013; Sanfilippo et al. 2016). Generation of a complete CA4-A genomic island required three steps: i) a translocation of *unk10* into the proto-CA4 genomic island, ii) acquisition of *fciC*, a putative ribbon helix-helix domain-containing regulator that has similarity to bacterial and phage repressors (Humily et al. 2013), and iii) acquisition of *mpeZ*, possibly by duplication and divergence of *mpeY*, then translocation into the proto-CA4-A genomic island (Fig. 6). It would also require the acquisition of the proper regulatory elements that are still unidentified.

Creation of the PT 3c-type PBS rod region from the same precursor that led to PT 3dA required three events: i) the loss of *mpeV*, ii) the translocation of *unk10* between *mpeU* and *cpeZ*, and iii) the divergence of the pre-existing lyase gene *mpeY* to make the lyase-isomerase gene *mpeQ* (Grébert et al. 2021). Then, the development of the PT 3dB from a PT 3c precursor only required the incorporation of a CA4-B genomic island in which the *mpeW* gene likely originated, like *mpeZ*, from duplication and divergence of *mpeY*, then translocation into the genomic island.

As previously noted, PTs 3c and 3dB share the same alleles for all PBS genes. Their only difference is the insertion of the CA4-B genomic island within the PBS rod region. Thus, conversion between these two PTs appears to be relatively straightforward and may occur frequently. In contrast, the PT 3a and 3dA PBS-encoding regions differ by a number of genes and often have different alleles for orthologous genes. Thus, although the acquisition of a CA4-A island theoretically should be sufficient to transform a PT 3a-type green light specialist into a chromatic acclimater (Sanfilippo, Nguyen, et al. 2019; Sanfilippo, Garczarek, et al. 2019; Grébert et al. 2021), the divergence between the PT 3a and 3dA PBS components could make this conversion problematic. In accordance with this, although a number of PT 3a strains have naturally acquired either a complete (MVIR-18-1) or a partial (WH8016 and KORDI-49) CA4-A genomic island (so-called PT 3aA strains), none exhibit a functional CA4 phenotype (Choi and Noh 2009; Humily et al. 2013).

### Lateral gene transfer did not occur at high frequency during the evolution of PBS

The inconsistency between phylogenies obtained from PBS and core genes has led several authors to suggest that frequent lateral gene transfer (LGT) events of parts of or the whole PBS rod region likely occurred during the evolution of *Synechococcus* (Six et al. 2007; Dufresne et al. 2008; Haverkamp et al. 2008; Everroad and Wood 2012; Sánchez-Baracaldo et al. 2019). However, by examining additional representatives of each PT/clade combination in this manuscript, we have shown that different alleles of the PBS genes correspond to the different PTs, and that the evolutionary history of each of these alleles is finely structured and globally consistent with the core phylogeny (Fig. 4). This suggests that LGT events between distantly related lineages are actually rare. An unambiguous LGT event occurred in strain MEDNS5 (PT 3c, clade VIa), since its *mpeBA* sequence clustered with PT 3c/clade III *mpeBA* sequences (Fig. 4 and Table S6). Similar observations were made for other PBS genes such as *cpeBA, mpeW/Y/Z* and *mpeU* (Humily et al. 2013; Mahmoud et al. 2017; Grébert et al. 2018). This suggests that there have been transfers of blocks of co-functioning genes. The match between the evolutionary history of each allele and the corresponding core phylogeny also suggests that most transfer events occurred very early during the diversification of marine *Synechococcus*. Indeed, the reconciliation analysis using ALE detected a high frequency of transfers to very ancient lineages in *Synechococcus* phylogeny (Fig. S10 and Table S6). However, the reconciliation model implemented in ALE does not account for incongruences between gene trees and species trees arising when an ancestral polymorphism in a population —in the present case, occurrence of several alleles— is not fully sorted (i.e., resolved into monophyletic lineages) after a speciation event. This is because of the stochastic way in which lineages inherit alleles during speciation (Tajima 1983; Galtier and Daubin 2008; Lassalle et al. 2015), and such incongruences are interpreted by ALE as replacement transfers (i.e., a transfer and a loss). This phenomenon, called ‘incomplete lineage sorting’ (ILS), is predicted to occur for at least some genes in a genome and is expected to be particularly important in prokaryotes with large population sizes (Retchless and Lawrence 2007; Degnan and Rosenberg 2009; Retchless and Lawrence 2010). In fact, the coalescent time (i.e. time to the last common ancestor), and hence the frequency of ILS, is predicted to be proportional to the effective population size (Abby and Daubin 2007; Batut et al. 2014). Since *Synechococcus* is the second most abundant photosynthetic organism in the oceans (Flombaum et al. 2013), we can reasonably expect to observe some ILS in this lineage. Assuming that the *Synechococcus* effective population size is the same order of magnitude as that estimated for *Prochlorococcus* (10^11^ cells; Baumdicker et al. 2012; Batut et al. 2014; Kashtan et al. 2014) and that *Synechococcus* has a generation time of about one day, a tentative allelic fixation time would be of about 280 million years (My). This rough estimate is on the same order of timescale as the divergence between SC 5.3 and SC 5.2/SC 5.1 (400-880 My ago) or, within SC 5.1, between marine *Synechococcus* and *Prochlorococcus* (270-620 My ago; Sánchez-Baracaldo et al. 2019). This further supports the possibility of ILS being the major source of the apparent incongruence in PT distribution between clades (Retchless and Lawrence 2007). This new evolutionary scenario would imply that the different PTs appeared before the diversification of SC 5.1 clades, and very likely before the divergence of SC 5.1 and 5.3, as was also recently suggested (Sánchez-Baracaldo et al. 2019). The basal position of the two SC 5.3 isolates in the phylogeny of the different *mpeBA* alleles (Fig. 4) reinforces this hypothesis. In this view, *Synechococcus* clades were derived from an ancestral population in which all PT 3 (a through f) co-existed. Some clades seem to have lost some PTs in the course of their separation from other clades such as clade IV or CRD1, in which we only observe isolates of PT 3dA. Others might have conserved most pigment types, such as clade II, which encompasses all PT 3 except 3dA. Thus, recombination would maintain intra-clade diversity (Lassalle et al. 2015), while allowing clades to expand to novel niches defined by environmental parameters such as iron availability or phosphate concentration (Doré et al. 2020). This would have allowed the partial decoupling of adaptation to multiple environmental factors from adaptation to light color (Retchless and Lawrence 2007; Retchless and Lawrence 2010).

In conclusion, the analyses the PBS rod region of newly sequenced *Synechococcus* isolates and of those retrieved from wild populations allowed us to clarify previous findings regarding the relationships between gene content and organization of this region, allelic variability and *Synechococcus* PTs. We proposed a scenario for the evolution of the different PTs and present a new hypothesis based on population genetics to explain the observed discrepancies between PT and core phylogenies. These results demonstrate that analyzing *Synechococcus* evolution from the perspective of its demographic history provides a promising avenue for future studies.

## Materials and methods

### Genome information

Genomic regions used in this study were obtained from 69 public complete or draft genomes (Dufresne et al. 2008; Cubillos-Ruiz et al. 2017; Lee et al. 2019; Doré et al. 2020). Information about these genomes can be found in Table S1.

### Fosmid libraries

Samples for construction of the fosmid library were collected during oceanographic cruises CEFAS (North Sea), BOUM (Mediterranean Sea) and the RRS Discovery cruise 368 (northeastern Atlantic Ocean) as well as from three long-term observatory sites. Two belong to the “Service d’Observation en Milieu Littoral” (SOMLIT), Astan located 2.8 miles off Roscoff and ‘Point B’ at the entrance of the Villefranche-sur-mer Bay, while the ‘Boussole’ station is located 32 miles off Nice in the ligurian current (Fig. 2A; (Antoine et al. 2008). Details on the sampling conditions, dates and locations are provided in Table S3. Pyrosequencing of the *petB* gene, cell sorting, DNA extraction, whole genome amplification, fosmid library construction, screening and sequencing were performed as previously described (Humily et al. 2014) and the fosmid libraries previously obtained from the Astan station were re-assembled using a different approach, as described below.

Sequencing reads were processed using BioPython v.1.65 (Cock et al. 2009) to trim bases with a quality score below 20, after which reads shorter than 240 nt or with a mean quality score below 27 were discarded. Reads corresponding to the fosmid vector, the *E. coli* host or contaminants were removed using a BioPython implementation of NCBI VecScreen (https://www.ncbi.nlm.nih.gov/tools/vecscreen/about/). Paired-reads were merged using FLASH v1.2.11 (Magoč and Salzberg 2011), and merged and non-merged remaining reads were assembled using the CLC AssemblyCell software (CLCBio, Prismet, Denmark). Resulting contigs were scaffolded using SSPACE v3.0 (Boetzer et al. 2011), and scaffolds shorter than 500 bp or with a sequencing coverage below 100x were removed. To reduce the number of contigs while preserving the genetic diversity, a second round of scaffolding was done using Geneious v6.1.8 (Biomatters, Auckland, New Zealand). Assembly statistics are reported in Table S4. Assembled scaffolds were manually examined to control for obvious WGA-induced as well as assembly chimeras. Annotation of PBS genes was performed manually using Geneious and the Cyanorak v2.0 information system (http://application.sb-roscoff.fr/cyanorak/). Plotting of regions was conducted using BioPython (Cock et al. 2009).

### Phylogenetic analyses

Sequences were aligned using MAFFT v7.299b with the G-INS-i algorithm (default parameters; Katoh and Standley 2013). ML phylogenies were reconstructed using PhyML v20120412 using both SPR and NNI moves (Guindon and Gascuel 2003). Phylogenetic trees were plotted using Python and the ETE Toolkit (Huerta-Cepas et al. 2016).

### Inference of evolutionary events

Species (vertical) phylogeny was inferred from a set of 73 conserved marker genes (Table S5; (Wu et al. 2013). For each marker gene, protein sequences were extracted from 69 isolate genomes and 33 *Synechococcus* SAGs (Table S1; Berube et al. 2018), aligned using MAFFT, and the alignment trimmed using trimAl (Capella-Gutiérrez et al. 2009). Alignments were concatenated into a multiple alignment which was used for reconstruction of the species tree. Phylogenetic reconstruction was done using RAxML 8.2.9, with 100 searches starting from randomized maximum parsimony trees and 100 searches from fully random trees (Stamatakis 2014). Best tree was selected and 200 bootstraps computed. Next, gene trees were inferred for every gene present in more than half of the considered genomes. For each gene, protein sequences were extracted from genomes and aligned using MAFFT. The resulting alignment was used for phylogenetic reconstruction with RAxML, and 100 bootstraps computed. Evolutionary events (gene duplication, transfer, loss or speciation) were inferred for each gene from these bootstraps using ALE v0.4, which uses a maximum-likelihood framework to reconcile gene trees with the species tree (Szöllősi et al. 2013).

## Supporting information

Grebert et al_Supplemental Figures S1-S11

Grebert et al_Supplemental Tables S1-S6

## Acknowledgments

This work was supported by the collaborative program METASYN with the Genoscope, the French “Agence Nationale de la Recherche” programs CINNAMON (ANR-17-CE02-0014-01) and EFFICACY (ANR-19-CE02-0019) as well as the European Union program Assemble+ to F.P and L.G. and by National Science Foundation Grants (U.S.A.) MCB-1029414 and MCB-1818187 to D.M.K. We thank Fabienne Rigaud-Jalabert for collecting sea water, Thierry Cariou for providing physico-chemical parameters from the SOMLIT-Astan station and Thomas Hackl for useful discussions about tycheposons. We are also most grateful to the Biogenouest genomics core facility in Rennes (France) for *petB* sequencing and the platform of the Centre National de Ressources Génomiques Végétales in Toulouse (France) for fosmid library screening. We also warmly thank the Roscoff Culture Collection for maintaining and isolating some of the *Synechococcus* strains used in this study as well as the ABIMS Platform (Station Biologique de Roscoff) for help in setting up the genome database used in this study and for providing storage and calculation facilities for bioinformatics analyses.

