## Supplementary material for "Diversity and evolution of pigment types and the phycobilisome rod gene region of marine *Synechococcus* cyanobacteria": Grebert et al_Supplemental Figures S1-S11

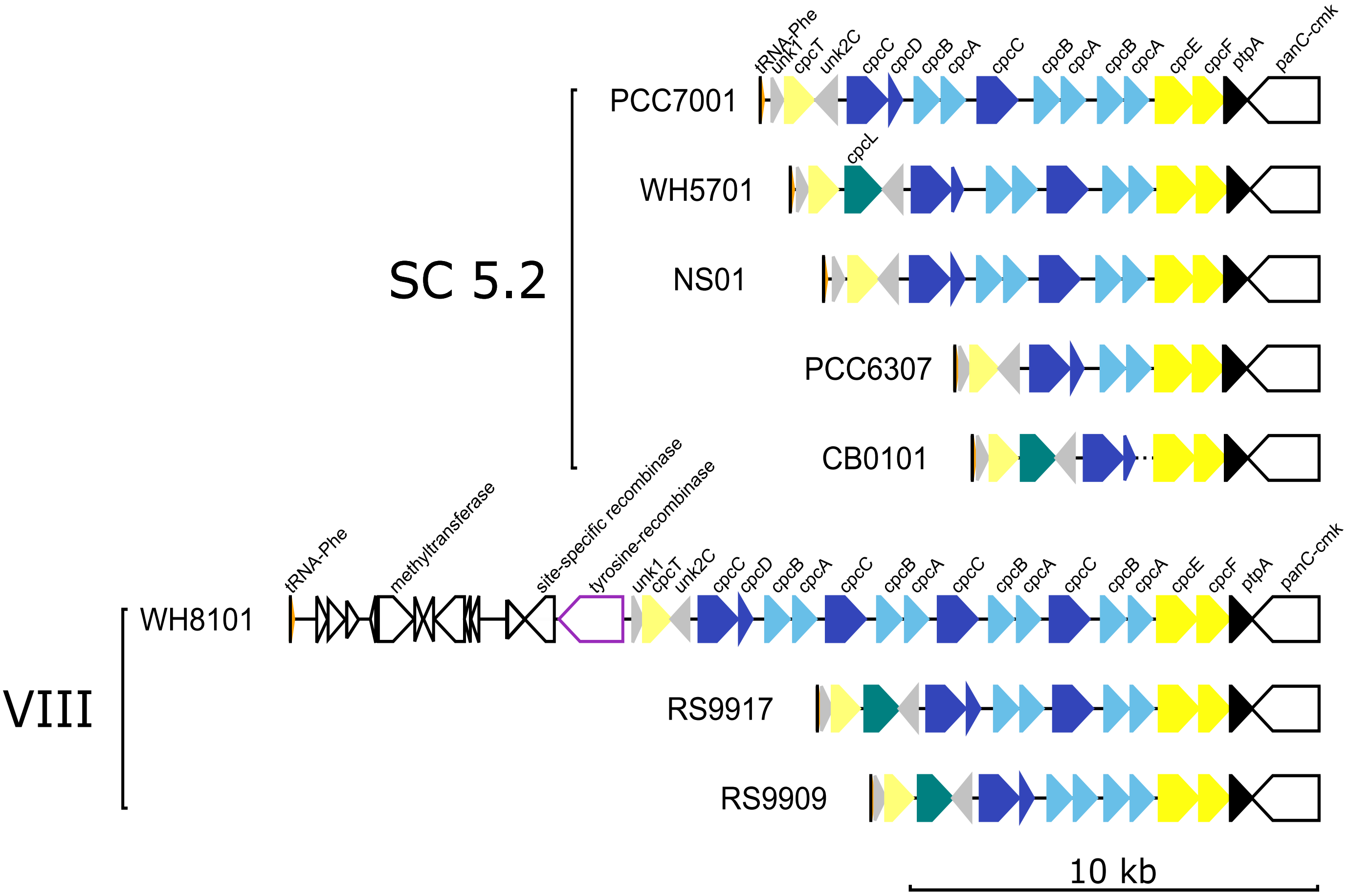


**Fig. S1: PBS rod region for all strains of pigment type 1.**
Regions are oriented from the phenylalanine tRNA to the conserved tyrosine phosphatase gene *ptpA*. PBS genes are coloured according to their inferred function (see insert in Fig. 1) and their length is proportional to the gene size. Strain CB0101 lacks *cpcBA* genes due to a gap in the genome assembly between *cpcC* and *cpcE*.

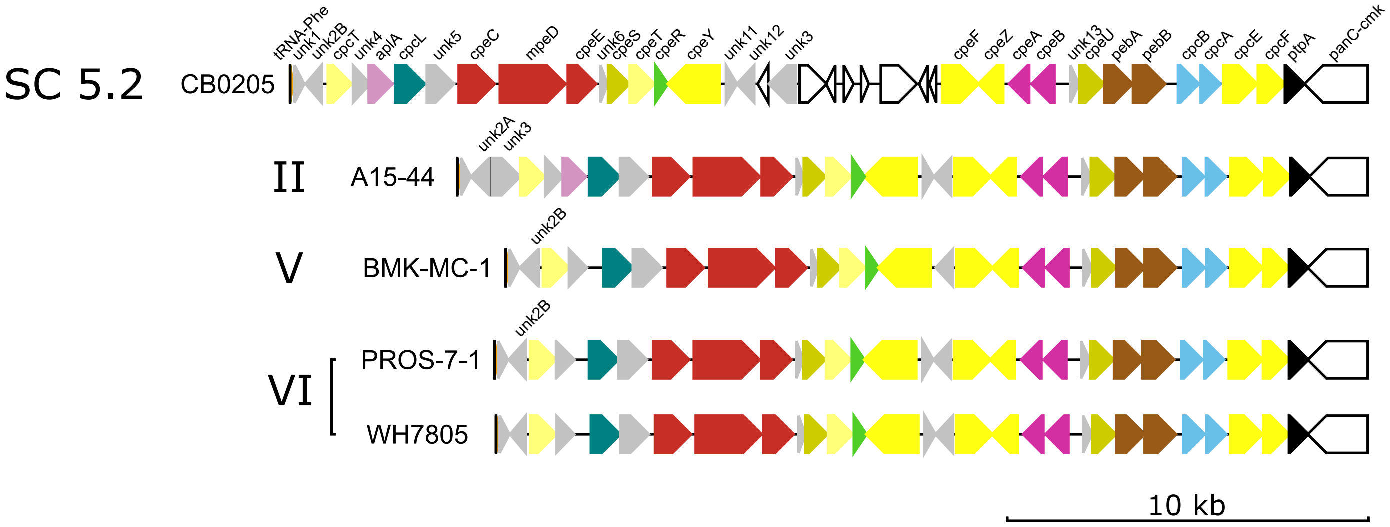


**Fig. S2: Same as Fig. S1 but for pigment type 2.**

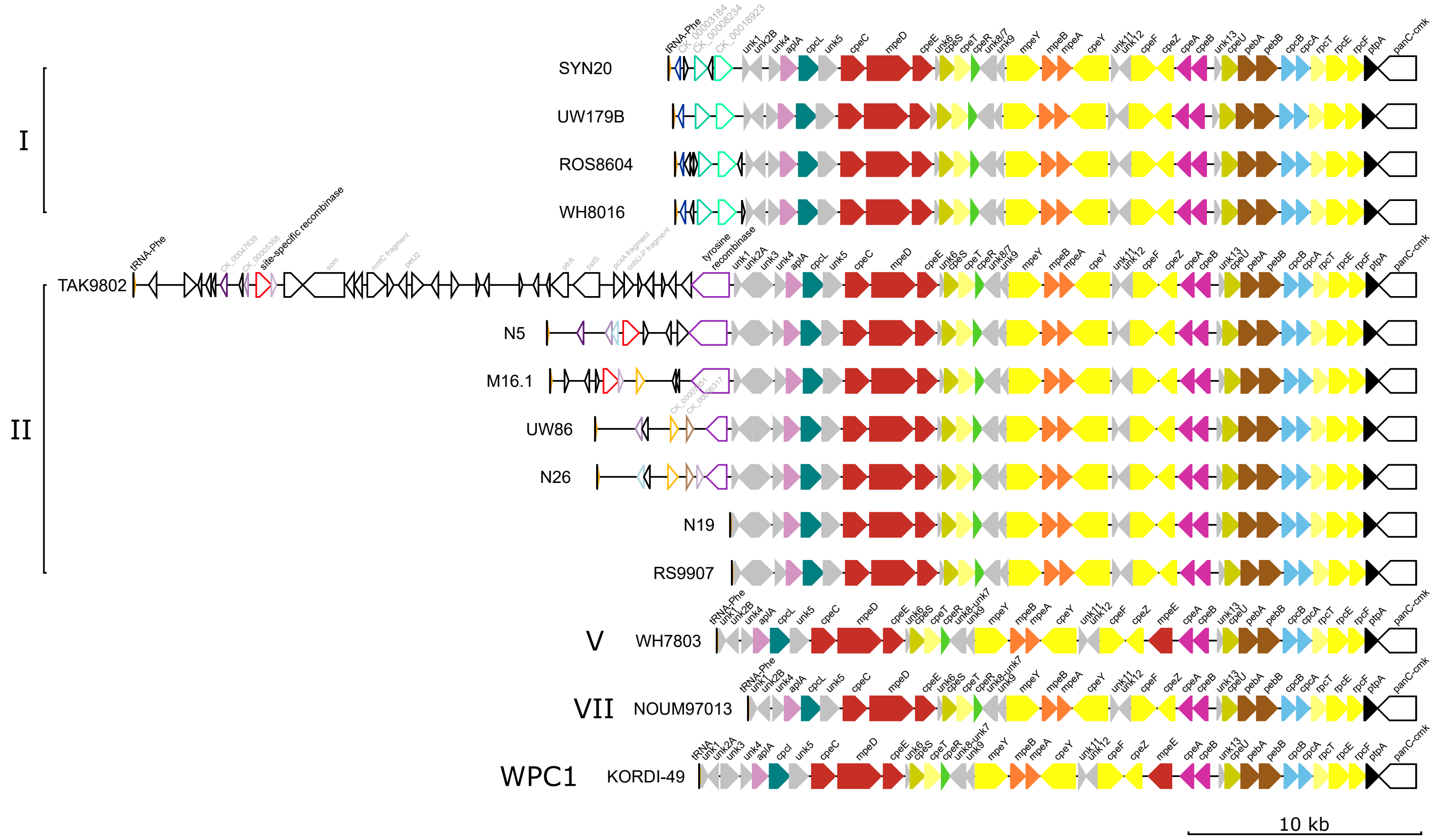


**Fig. S3: Same as Fig. S1 but for pigment type 3a.**

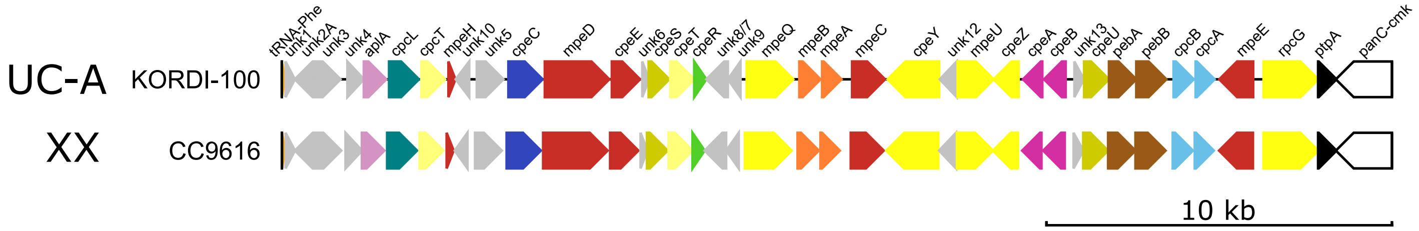


**Fig. S4: Same as Fig. S1 but for pigment type 3f.**

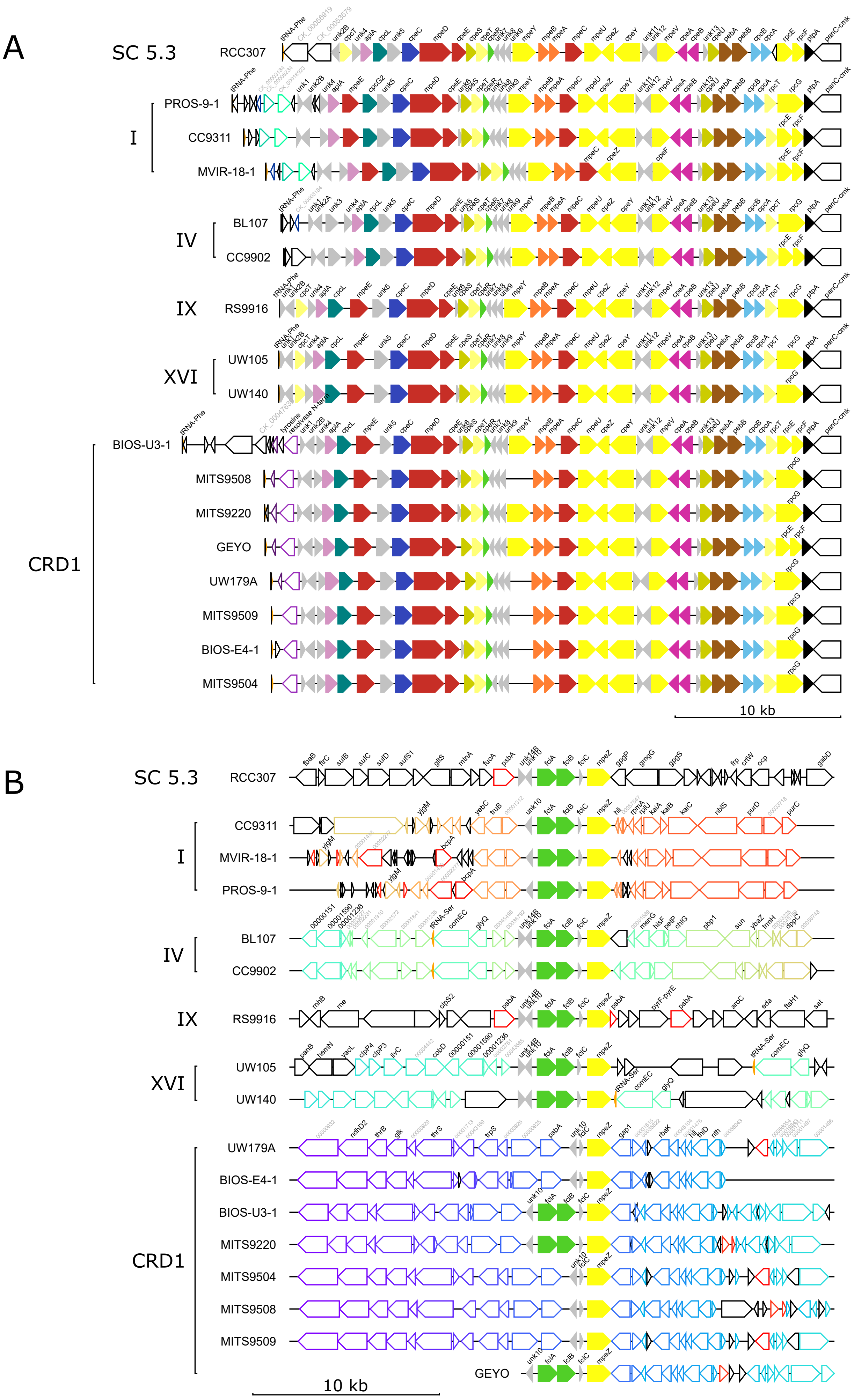


**Fig. S5: PBS rod region and CA4-A island for all strains of pigment type 3dA.**
(*A*) PBS rod region. (*B*) CA4-A genomic island and surrounding genes. Solid arrows represent genes from the CA4-A genomic island and are coloured according to their inferred function (see insert in Fig. 1). Open arrows are neighbouring genes, and orthologs in different strains are shown with the same contour color.

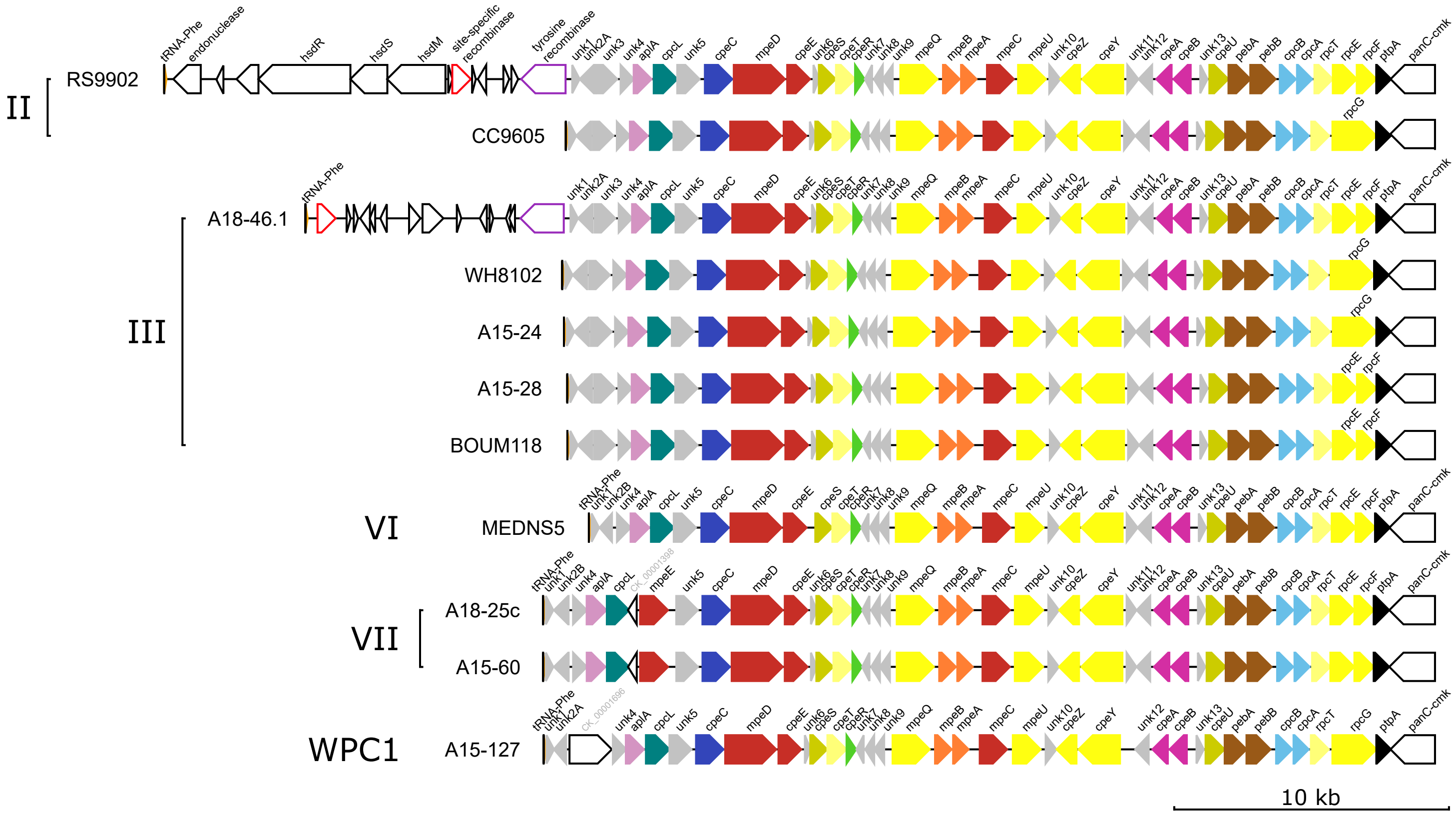


**Fig. S6: Same as Fig. S1 but for pigment type 3c.**

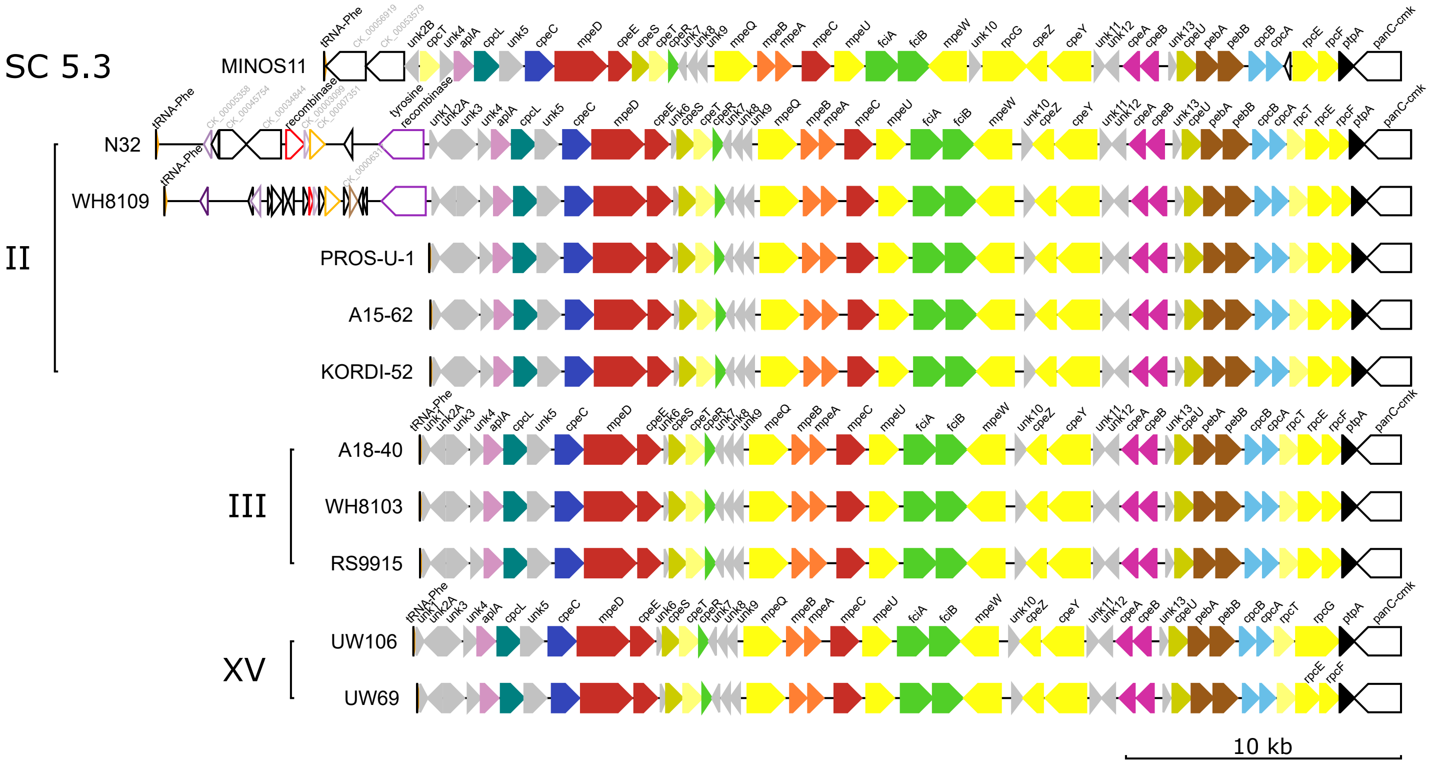


**Fig. S7: Same as Fig. S1 but for pigment type 3dB.**

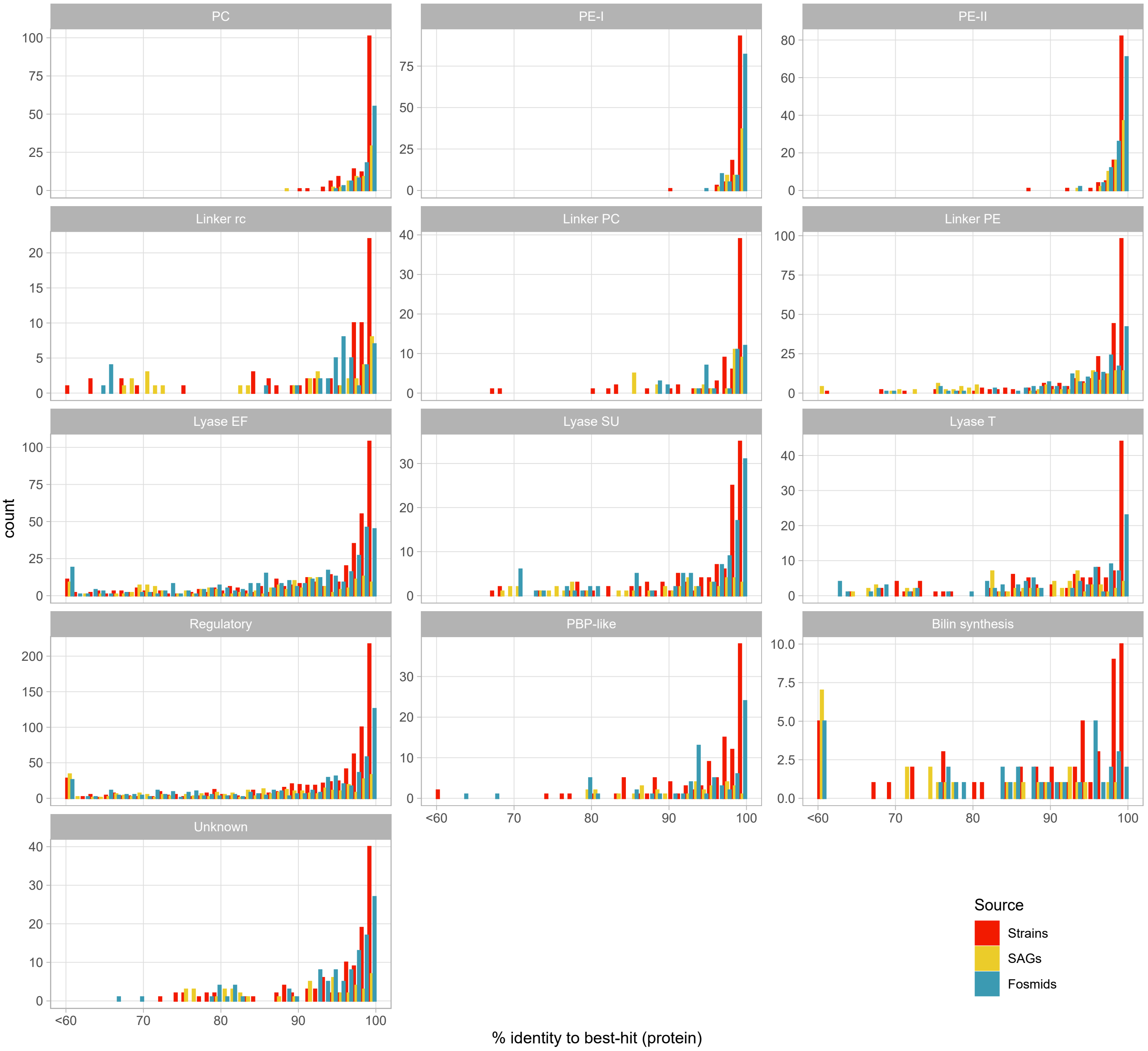


**Fig. S8: Genetic variability of the different proteins encoded in the PBS rod genomic region, grouped by functional categories.** For each group of orthologous proteins, identity is based on the blastp best-hit other than self-hit for strains, and blastp best-hit to all strains for fosmids and SAGs, and results for each group of orthologs have then be gathered by functional categories as follows: PC, phycocyanin (CpcA/B, RpcA/B); PE-I, phycoerythrin-I (CpeA/B); PE-II, phycoerythrin-II (MpeA/B); Linker rc, linker rod-core-like (CpcL); Linker PC, PC-associated linker (CpcC, CpcD, CpeC); Linker PE, PE-associated linker (CpeE, MpeC, MpeD, MpeE, MpeH); lyase EF, lyase of the E/F clan (CpeF, CpeY, CpeZ, MpeU, MpeY, MpeW, MpeZ, RpcE, RpcF, RpcG, CpcE, CpcF); lyase SU, lyase of the S/U clan (CpeS, CpeU); lyase T, lyase of the T clan (CpcT, CpeT, RpcT); Unknown, uncharacterized conserved hypothetical proteins (Unk1 through 13, Unk2A, Unk2B, Unk2C, Unk8/7, Unk14B).

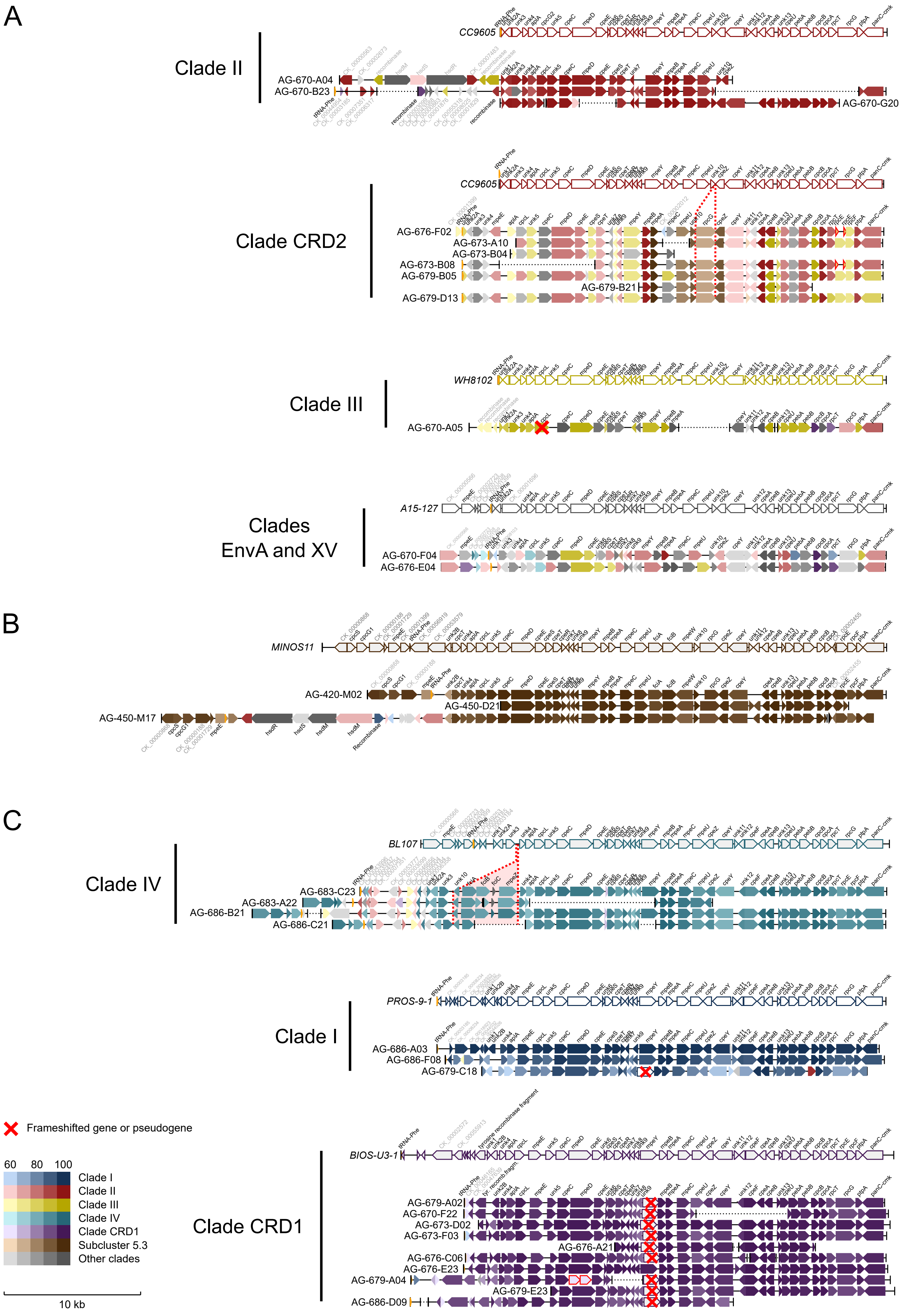


**Fig. S9: partial or complete PBS rod region retrieved from single-cell amplified genomes (SAGs).** PBS regions are grouped by pigment type, with PT 3c in (*A*), PT 3dB in (*B*) and PT 3dA in (*C*). Colors represent the clade of the reference strain giving the best blastp hit within the given pigment type.

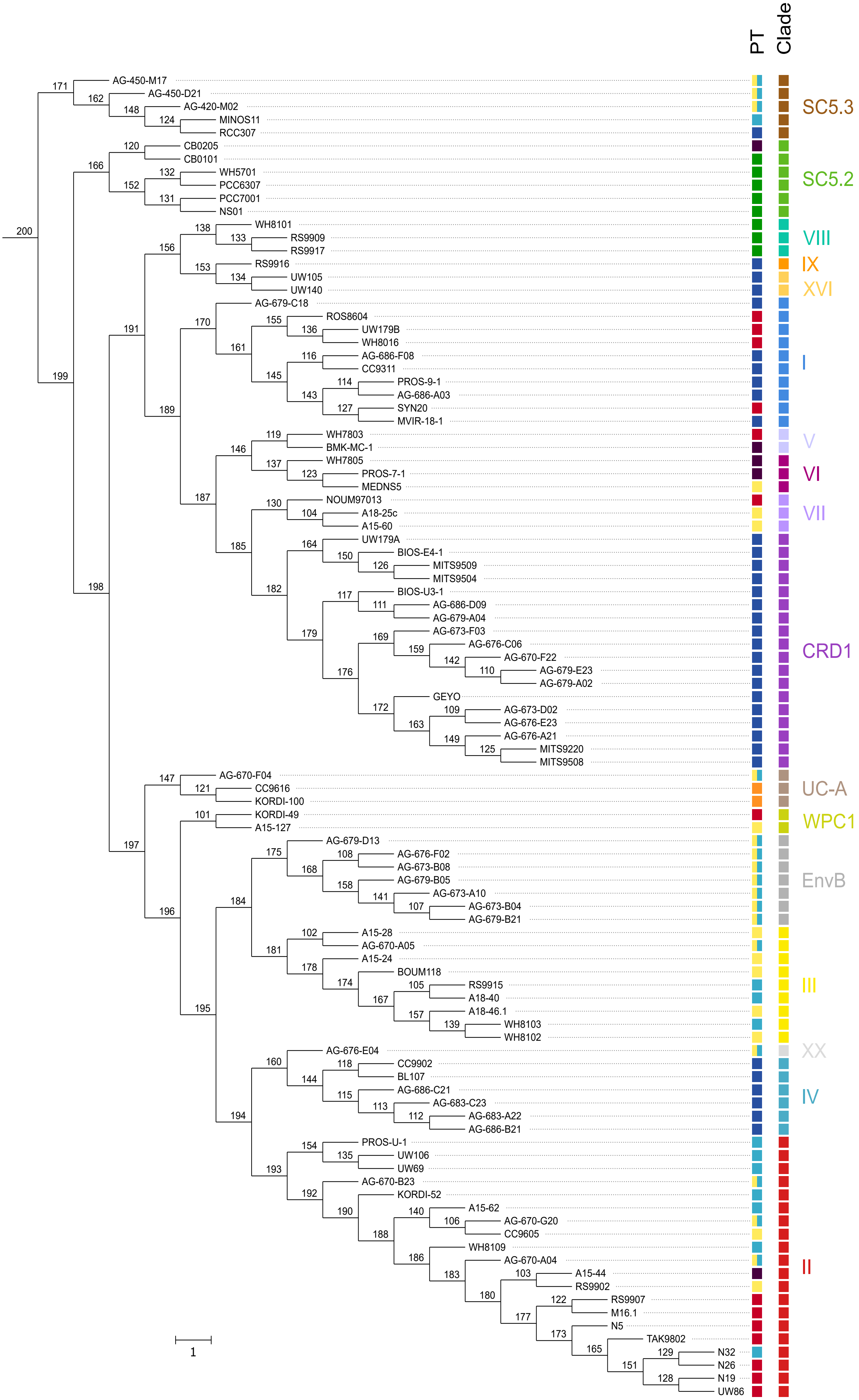


**Fig. S10: Reference species tree based on 73 core proteins used in ALE analysis.** All internal nodes are labelled according to ALE numbering, allowing identification of transfer events from/to internal nodes.

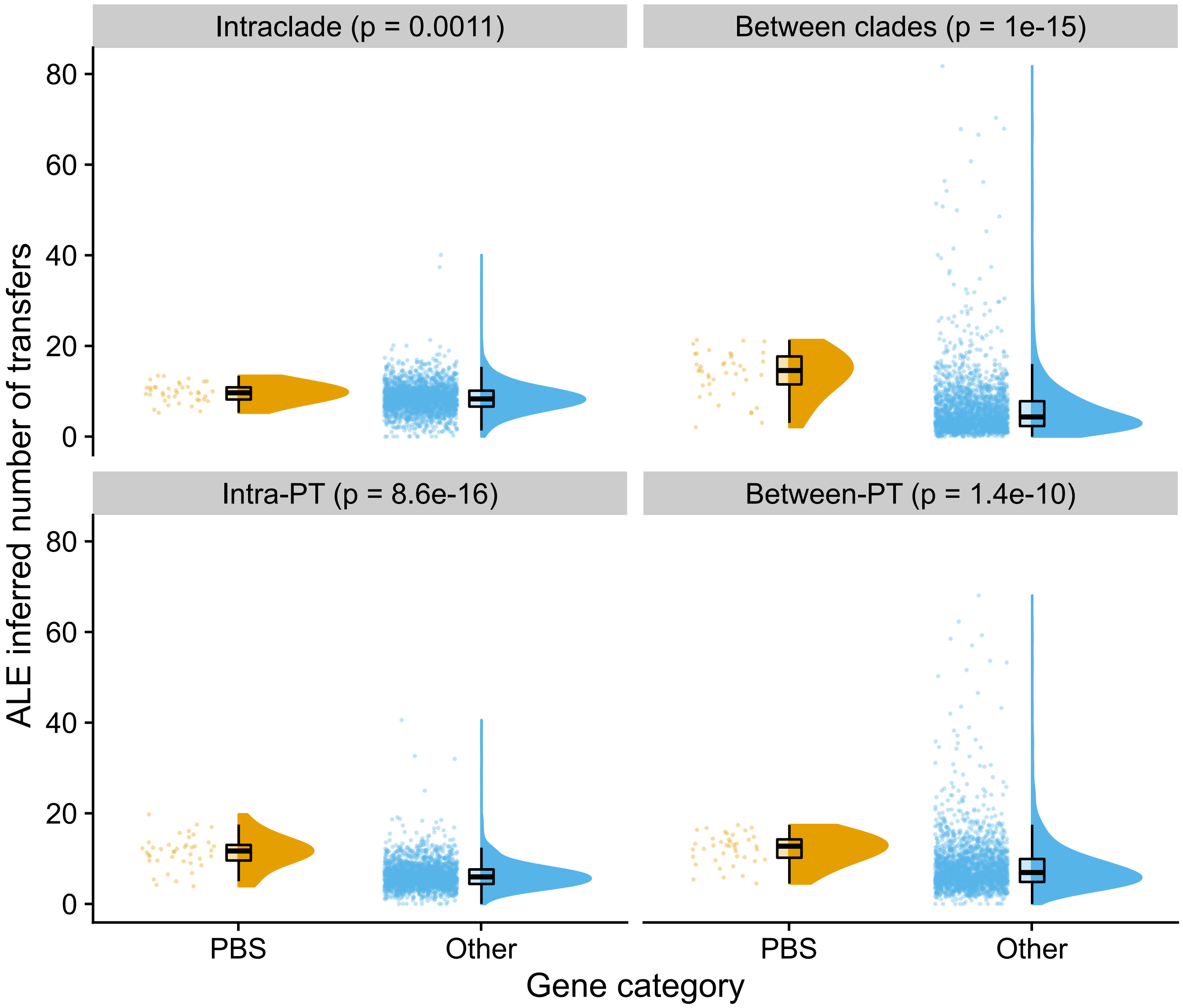


**Fig. S11: Comparison of the distribution of transfer events inferred by ALE for gene belonging to the PBS rod region (PBS) and other genes (Other).**

Transfer events were classified as intraclade if they involved two strains or ancestral lineages from the same clade, and between clades otherwise. Similarly, transfers were classified as intra-pigment type (Intra-PT) if they involved two strains or ancestral lineages having the same pigment type, and between pigment types (Between-PT) otherwise. P-values for Wilcoxon rank sum exact test are reported.
